# Cell-autonomous Wnt activity promotes transient re-programming and cell cycle re-entry of coronary artery endothelial cells

**DOI:** 10.64898/2026.02.23.707374

**Authors:** Bhavnesh Bishnoi, Alfia Nirguni Saini, Vinay Rao, Omkar Golatkar, Ravindra Kailasrao Zirmire, Shruthi Viswanath, Perundurai Subramaniam Dhandapany, Soumyashree Das

## Abstract

Coronary occlusion leads to formation of collateral arteries connecting occluded and perfused arteries. This process, namely artery reassembly, is temporally restricted to the cardiac regenerative window in mice and relies on cell cycle re-entry of post-mitotic artery endothelial cells (ECs). Analyses of single cell RNA sequencing data of ischemic mouse hearts led to identification of Wnt activity in proliferating artery cells. Using mouse genetics and whole organ imaging at cellular resolution, we show that Wntless (Wls)-mediated Wnt secretion and Frizzled (Fzd)4-dependent Wnt reception by artery cells, was essential for development of coronary collaterals. Specifically, Fzd4 promoted proliferation and expression of (progenitor-like) capillary/venous markers in coronary artery ECs. In embryonic hearts, coronary arteries originate from capillaries. Fzd4 in these capillaries was essential for proliferation; depletion of which, accelerated their differentiation into arteries by arterialization. Along these lines, an exogeneous dose of human WNT2-sFRP1 ligand complex reactivated artery cell proliferation and restored artery reassembly in non-regenerative mouse hearts. Thus, ischemia-induced cell-autonomous Wnt signaling promoted developmental reprogramming of coronary artery ECs. Finally, exome sequencing of coronary artery disease patients, revealed unique pathogenic variants of *WLS* and *FZD4*. Together, our study provides mechanistic insights into EC plasticity and underscores the significance of WNT pathway in cardiovascular disease.

## Introduction

Endothelial fate decisions made at the right time and place are essential for organ growth during development^1^ and tissue remodeling after injury^2,3^. Endothelial cells (ECs) arise *de novo* from the mesoderm and subsequently commit to arterial or venous fates to enable vascular expansion and patterning^4,5,6^. Later, an immature vascular plexus, of venous origin^7,8,9,10^, gives rise to arteries, veins, and capillaries, within organs. In heart, ECs of this initial vascular plexus exit cell cycle and differentiate into pre-artery ECs^7,11^. These pre-artery cells migrate and coalesce to form coronary artery that connects to, and receives blood flow from the aorta. Thus, ECs commit to an artery fate before the onset of arterial blood flow^12,13^. Subsequently, coronary pre-artery ECs expand by migrating against the direction of blood flow to contribute to the distal ends of arteries^11,14^. A similar mechanism operates in the retina, where venous ECs differentiate into artery ECs^15^, migrate against blood flow and contribute to the extension of distal artery tips^10^. Thus, inward migration of ECs, predominantly of venous origin and nature, directs organ-specific artery growth^16^.

The subsequent organ-specific vascular growth, during development or post-injury, heavily relies on capillary/venous ECs^10,17,18^. For the longest period, artery ECs were thought to be terminally differentiated. However, several recent studies have elaborated on context-dependent artery cell plasticity during development^19^, repair^20^ and regeneration^21,22,23^. For instance, following myocardial infarction (MI), arterial ECs dissociate from pre-existing coronary arteries as single cells, de-differentiate, proliferate, and migrate into the watershed region—the artery-free zone between the terminal branches of the left and right coronary arteries—to form collateral arteries. These collaterals connect the non-perfused ligated artery to a healthy artery, thereby alleviating ischemia and providing an alternative route for blood flow. This process of forming collateral arteries from pre-existing arteries, namely artery reassembly, is temporally restricted to the first seven days after birth in mice (called cardiac regenerative period)^21,22,24^. The precise genetic program, sequence of cellular events and the molecular trigger dictating arterial identities during artery reassembly, largely remains unknown.

Wnt (Wingless-related integration site) signaling drives artery fate^25,26^. Constitutive activation of canonical Wnt signaling downstream effector, β-catenin, promotes arterial identity during growth of embryonic^25^ and postnatal vasculature^27^. Similarly, deletion of Wnt agonist *R-spondin3* from cardiomyocytes^28^ or its cognate receptor *Lgr5* from endothelial cells^29^, impairs coronary artery development. The principal canonical Wnt receptor, Frizzled-4 (Fzd4), is expressed in both artery and capillary ECs, and regulates angiogenesis in retina^30^, inner ear, central nervous system^31^ and heart^32^. In mice, several Wnt ligands are expressed by cardiac cells^33^ which are upregulated post-MI^34^. Specifically, overexpression of canonical Wnt ligand 10b in cardiomyocytes increases arterial density post-MI^35^. Thus, Wnt pathway is instrumental in shaping the vascular growth in developing and regenerating hearts. However, given the wide distribution of its ligands and co/receptors across different cardiac cell-types, it is not well-understood how multiple Wnt signals from different cell types converge to generate a unifying EC response.

In this study, we investigate endothelial specific role of Wnt pathway during cardiac development and regeneration. We show that Fzd4 regulates artery cell specification by carefully balancing endothelial proliferation and differentiation, and more importantly, in a cell-autonomous manner. Interestingly, the inability to execute artery reassembly (by non-regenerative hearts) is overcome with an exogeneous dose of human WNT2–sFRP1 complex. Finally, identification of deleterious genetic variants of WNT pathway genes in heart patients of South (Asian) Indian origin supports the significance of our findings in mice, and underscores the importance of optimal Wnt activity in cardiac health. Together, we establish the role of WNT pathway in regulation of endothelial fate decisions associated with coronary artery and collateral development; processes key to a heathy heart.

## Results

### Artery reassembly depends on artery cells re-entering cell cycle

Proliferation of artery ECs is a key step to formation of coronary collateral arteries via artery reassembly (Figure **S1A**)^22^. To assess the role of cell-cycle re-entry of coronary artery cells, in artery reassembly, a cyclin dependent kinase (Cdk)-1 (responsible for G2 to M transition) (Figure **S1B**) was deleted from pre-existing arteries, prior to myocardial infarction (MI). No significant differences were observed in neonatal coronaries without arterial *Cdk1* (Figure **S2**). MIs were performed in mice with *Cx40CreER* and *Rosa26^TdTomato^*, where Connexin40^+^ (Cx40) arterial ECs are marked by TdTomato fluorescence. TdTomato signal is spatially restricted to *Cx40CreER*-expressing artery ECs and temporally controlled by Tamoxifen induction which activates the Cre recombinase. Resident neonatal coronary arteries were genetically labelled at postnatal day (P)0 or 1 through P2/3, when MI was performed (Figure **S1C**). Hearts were explanted 4 days post-MI and assessed for artery reassembly by quantification of TdTomato^+^ collaterals (Figure **S1C**). Since the Tamoxifen persists for 2 days^21,36^, any new TdTomato^+^ arteries appearing after this period would originate from pre-existing artery ECs and are lineage-traced with TdTomato. We first evaluated if proliferation was affected in single artery cells deleted for *Cdk1*. Hearts injected with EdU (5-ethynyl-2’-deoxyuridine), at the time of explant, were assessed for (EdU^+^ TdTomato^+^) lineage-traced proliferating artery ECs (Figure **S1C**). Our earlier studies show that ∼10% of single artery cells participating in artery reassembly are EdU^+ 21,22^. However, arterial deletion of *Cdk1* led to a ∼2-fold decrease (5.5% per heart) in proliferating artery ECs (Figure **S1D**), suggesting defective proliferation in absence of Cdk1. Deletion of *Cdk1* from pre-existing artery ECs led to a significant reduction in TdTomato^+^ collateral arteries between perfused (right or septal) and occluded (left) branches of coronary arteries. While control hearts showed ∼8 TdTomato^+^ collaterals per heart, arterial knockouts for *Cdk1* showed ∼4 collaterals per heart (Figure **S1E**). This was also evident from images of whole hearts and ischemic watersheds from *Cdk1^L/L^; Cx40CreER*; *Rosa26^TdTomato^* mice, which show scarcity of TdTomato^+^ single artery cells and collaterals, as compared to their control counterparts (Figure **S1F**-**I**, compare **G** to **I**). Together, Cdk1-dependent cell-cycle re-entry of artery ECs is essential for artery reassembly.

### Secreted Wnt from artery ECs promotes proliferation upon MI

To identify potential signalling pathway regulating artery EC proliferation, in response to MI, single cell RNA sequencing data from neonatal and adult mouse hearts generated by Arolkar et al, was reanalysed (Figure **S3A**)^22^. The neonatal dataset was generated from Cx40^+^ lineage labelled/traced artery ECs, 2-3 days post-MI (Figure **S3A**). Endothelial cells from regenerative neonatal hearts capable of artery reassembly were subjected to PROGENy (Pathway RespOnsive GENes for Activity Inference) and assessed for pathway activities (Figure **S3B**). Specifically, three endothelial cell clusters expressing artery EC markers were analyzed; 1. proliferating artery ECs (cycling saECs), 2. mature artery ECs and 3. transient ECs, which expressed both capillary and artery markers^22^. Interestingly, Wnt and PI3K signalling were upregulated in cycling single artery ECs, but downregulated in mature and transient ECs (Figure **S3B**), suggesting a role for these molecules in post-MI artery cell proliferation.

Wnt pathway regulates coronary artery development^28^ and optimizes postnatal arterial density^32^. Hence, we investigated the role of Wnt pathway (Figure **S3C**) in artery reassembly─a regenerative event triggered only post-MI. Wnt ligand secretion is facilitated by a pan-Wnt ligand carrier protein, Wntless (Wls). Upon secretion, Wnt ligands bind to Frizzled receptors, and trigger a cascade of molecular events which stabilizes β-catenin, and transports it to nucleus. Nuclear β-catenin promotes transcription of molecules like *Axin2*, *Myc*, *Ccnd1*, which regulate cellular proliferation and fate decisions (Figure **S3C**). We first checked if downstream effectors of Wnt pathway like Axin2 and β-catenin are expressed by coronary artery ECs (Figure **S3D**-**G**). Cryosections of healthy P6 hearts from *Axin2d2EGFP* mice were immunolabelled for EGFP and artery EC specific marker, Cx40, (Figure **S3D**, **F**), to assess colocalization of EGFP (Axin2) with Cx40 expressing arteries. Similarly, cryosections of *Cx40CreER; Rosa26^TdTomato^* hearts were immunolabelled for β-catenin (Figure **S3E**, **G**) and assessed for colocalization of β-catenin and TdTomato (artery ECs). Axin2 (marked with antibody against GFP) and β-catenin, showed specific immunostaining in cardiac cells including coronary artery ECs (Figure **S3D**, **E**), with β-catenin prominently labelling cellular junctions (Figure **S3E**, insets). Together, Wnt downstream effector molecules such as Axin2 and β-catenin are present in mouse coronaries, postnatally.

Wntless (Wls) regulates secretion of Wnt ligands across tissues and species^37,38^. We analyzed the expression of *Wls* and *Wnt* ligands in our scRNA dataset^22^, pre- and post-MI (Figure **S4**). In Arolkar et al, post-MI datasets were generated at a timepoint prior to completion of artery reassembly^21^, i.e. 2/3 days post-MI in neonates, and 9/11 days post-MI in adults^22^ (Figure **S4A**, **B**). Clustering based on gene expression showed that mature and cycling artery ECs were common to neonatal and adult coronaries^22^ (Figure **S4C**, **E**). However, transient EC cluster was exclusive to neonates and capillary EC cluster was specific to adults (Figure **S4C**, **E**)^22^. *Wls* was expressed in all EC clusters in neonates (Figure **S4D**) and adults (Figure **S4F**). Neonatal and adult mouse coronary arteries expressed 14 (Figure **S4G**, **H**) and 13 (Figure **S4I**, **J**) *Wnt* ligands respectively, both in sham and MI hearts. Thus, *Wnt* ligands, and their carrier-protein *Wls*, are expressed in mouse coronaries post-MI.

Given the indispensable role of Wnt pathway components in shaping coronary artery network^28,32^, we first checked if active secretion of Wnt ligands impacts artery-reassembly. We depleted Wls from artery ECs. *Wls* floxed mice in conjunction with Tamoxifen inducible *Cx40CreER; Rosa26^TdTomato^* mice were used. Of note, postnatal hearts with knockout for *Wls*, specifically in artery ECs, did not show any gross morphological phenotype (Figure **S2**). Tamoxifen was injected at P0/P1 followed by MI at P2/P3 and hearts were analyzed for artery reassembly at P6/P7 (Figure **1A**), 4 days post-MI. Whole hearts were imaged at cellular resolution to visualize TdTomato^+^ lineage-traced collateral arteries formed by artery reassembly (Figure **1B**-**F**). Control hearts (Figure **1B**) showed an average of 10 collaterals in the watershed region, spanning between branches of ligated left coronary artery and healthy right coronary artery (Figure **1D**). Additionally, TdTomato^+^ single artery ECs were abundant in this collateral forming zone (Figure **1E**, arrowheads). In contrast to this, hearts with arterial knockout of *Wls* showed 2.5-fold reduction in TdTomato^+^ collateral arteries (Figure **1C**, **D**, **F**), despite the presence of single artery ECs in the watershed area (Figure **1F**, arrowheads and brackets). Thus, arterial Wls regulates collateral formation by artery reassembly.

**Figure 1:**
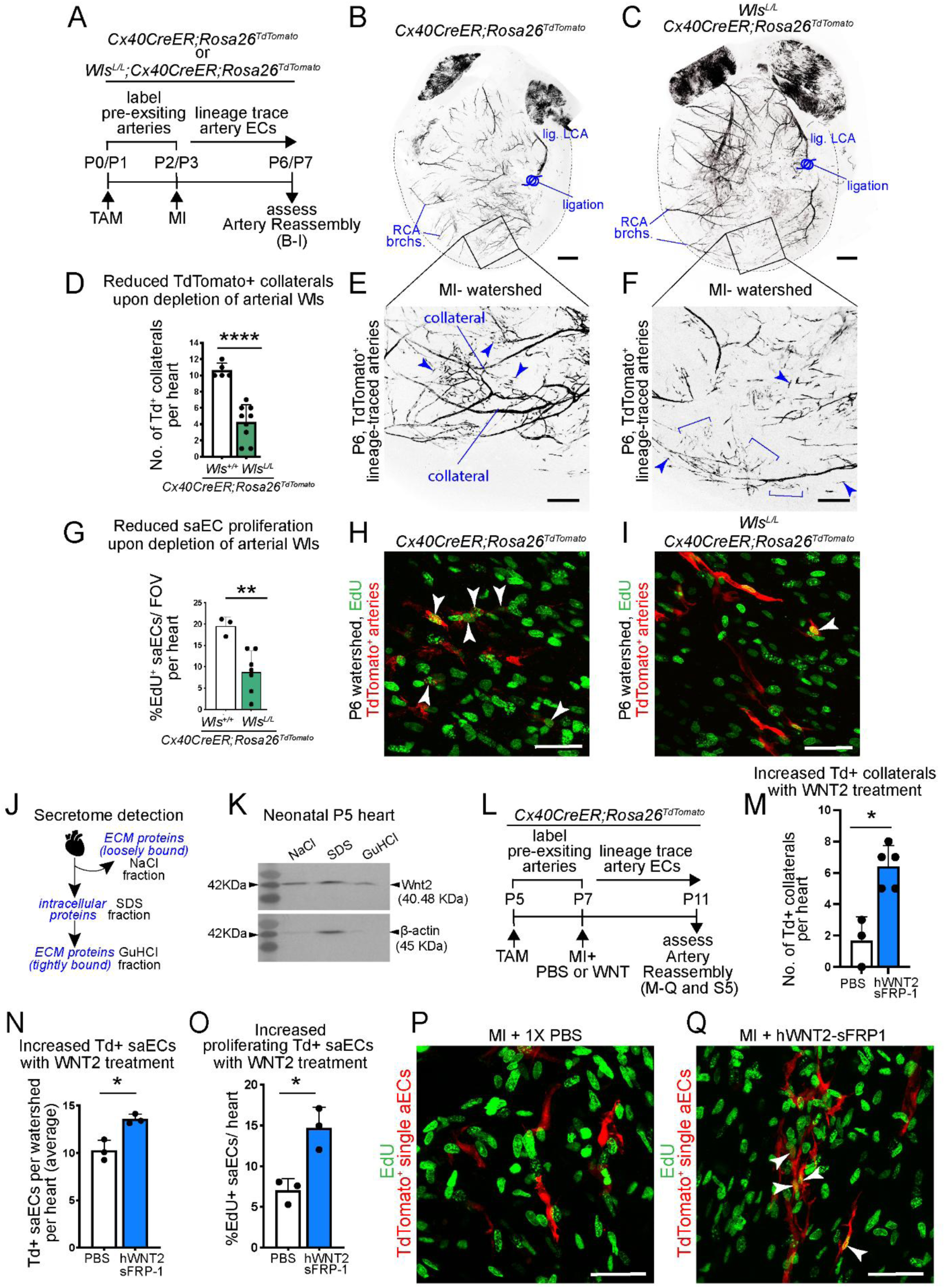
Wls-dependent Wnt secretion promotes artery reassembly. (**A**) Experimental design to assess the role of arterial Wls in artery reassembly. Confocal images of whole neonatal hearts from (**B**) control (*Cx40CreER; Rosa26^TdTomato^*) and (**C**) arterial knockouts for Wls (*Wls^L/L^; Cx40CreER; Rosa26^TdTomato^*). Black shows lineage traced arterial endothelial cells. (**D**) Quantification of TdTomato^+^ collaterals per heart. p<0.0001. (**E**, **F**) Watershed areas from (**E**) control (inset from **B**) and (**F**) arterial knockouts for *Wls* (inset from **C**). Black shows lineage traced arterial endothelial cells. Arrowheads and brackets point to single and cluster of lineage traced artery cells, respectively. (**G**) Quantification of EdU^+^ proliferating single artery cells post-MI. p=0.0013. (**H**, **I**) Representative confocal images of watershed regions from (**H**) control and (**I**) arterial knockouts for *Wls*, showing EdU^+^ proliferating cells in green and TdTomato^+^ lineage traced artery endothelial cells in red. Arrowheads point to EdU^+^ TdTomato^+^ proliferating single artery cells, post-MI. (**J**) Working principle underlying detection of proteins in cardiac secretome. (**K**) Western blot showing presence of Wnt2 in uninjured cardiac cells and secretome. β-actin protein is enriched intracellularly (in SDS fraction). Wnt2 was found intracellularly and associated with extracellular matrix. (**L**) Experimental design to assess artery reassembly upon injection of hWNT2-sFRP1 into non-regenerative hearts. (**M**) Quantification of TdTomato^+^ collaterals per heart when injected with 1X PBS or 55ng of hWNT2-sFRP1. p=0.0123. **N** shows quantification of total number of TdTomato^+^ single artery endothelial cells per watershed per heart. p=0.0199. **O** shows quantification of EdU^+^ TdTomato^+^ (proliferating) single artery cells found per watershed per heart. p=0.0171. (**P**, **Q**) show confocal images of P11 cardiac watershed area, injected with either (**P**) 1X PBS or (**Q**) hWNT2-sFRP1. TdTomato^+^ arteries are in red and EdU^+^ nuclei are in green. Arrowheads point to TdTomato^+^ single artery ECs with EdU^+^ nuclei. Note that TdTomato^+^ single artery ECs do not colocalize with EdU in **P**. Scale bars: **B**, **C**, 500µm; **E**, **F**, 200µm; **H**, **I**, 50µm; **P**, **Q**, 50µm. P, postnatal day; Tam, Tamoxifen; Td, TdTomato; MI, Myocardial Infarction; inj, injection; lig, ligation; LCA, left coronary artery; RCA, right coronary artery; saEC, single artery endothelial cell; h, human; sFRP1, soluble Frizzled Receptor Protein 1.

Wnt pathway is essential for proliferation and coronary artery development in mice^28,32^. We asked if *Wls* deleted TdTomato^+^ single artery ECs can proliferate in response to MI (Figure **1G**-**I**). To check this, neonates were injected with EdU, 4 days post-MI. EdU^+^ TdTomato^+^ proliferating lineage-traced single artery ECs were visualized using confocal microscopy (Figure **1H**, **I**) and quantified in control and *Wls* knockout hearts (Figure **1G**). In contrast to the controls, EdU^+^ single artery ECs showed 2.2-fold reduction in hearts depleted for arterial Wls (Figure **1G**). Thus, Wnt secreted by artery ECs promote their proliferation.

### WNT2-sFRP1 activates artery reassembly in non-regenerative neonatal hearts

Arterial expression of *Wnt2 and 9b* mRNA in regenerative neonatal hearts increases, post-MI (Figure **S4G**, **H**). This increase was not evident in adult artery EC cluster (Figure **S4I**, **J**), indicating a potential role of these proteins in artery reassembly. Wnt2 has a pronounced function in endothelial cell biology^39,40^. Hence, we first checked the presence of Wnt2 protein in neonatal hearts. We used a biochemical method that utilizes decellularized cardiac tissue to detect extracellular matrix (ECM) bound proteins^41^. ECM bound protein secretome were generated by subsequent incubations with NaCl, SDS and GuHCl. Distinct ECM fractions were then subjected to Wnt2 protein detection with western blotting. While SDS fraction contained intracellular proteins, NaCl and GuHCl fractions contained ECM bound proteins (Figure **1J**). As expected, a β-actin protein band was primarily observed in (intracellular) SDS fraction, though, a faint band was also observed in NaCl fraction (Figure **1K**). Interestingly, Wnt2 protein was observed in all three fractions suggesting that it is present intracellularly and, in an ECM-bound form in regenerative hearts.

Older non-regenerative hearts fail to execute artery reassembly^21^. In adult hearts, arterial expression of *Wnt2* mRNA did not increase post-MI (Figure **S4I**, **J**). We checked if an exogenous dose of Wnt2 protein could trigger artery reassembly in these hearts. *Cx40CreER; Rosa26^TdTomato^* mice were injected with Tamoxifen at P5 followed by MI at P7 with a simultaneous intra-cardiac injection of either 1XPBS or 55ng hWNT2-sFRP1 (human WNT2 conjugated to secreted Frizzled Related Protein)-1. Hearts were explanted and assessed for collateral formation by artery reassembly at P11, 4 days post-MI (Figure **1L**). When compared to PBS, hWNT2-sFRP1 injected P11 MI hearts triggered artery reassembly (Figure **1M**-**O** and Figure **S5**). Hearts receiving an exogenous dose of hWNT2-sFRP1 showed 4-fold increase in TdTomato^+^ collateral artery number as compared to the control counterparts (Figure **1M**, average collaterals in hearts treated with WNT2: 6.4, PBS: 1.6). Activation of artery reassembly program was also evident by a massive expansion of TdTomato^+^ arterial lineage in WNT2 treated hearts (Figure **1N** and **S5A, B**). Unlike PBS treatment (Figure **S5C**, **D**), watershed regions of P11 hearts which received WNT2 injection (Figure **S5E**-**G**), showed several TdTomato^+^ lineage-traced single (arrowheads in Figure **S5F**-**H**) or clustered artery cells (brackets in Figure **S5F**, **H**) populating the watershed regions. Additionally, TdTomato^+^ arterial conduits (including collateral arteries) were frequently found in WNT2 injected watershed regions (arrows in Figure **S5F**, **H**). Thus, the number of TdTomato^+^ single artery ECs were significantly increased in hearts treated with WNT2 as compared to PBS (Figure **1N**). Interestingly, EdU assays on both hearts, either treated with PBS or hWNT2-sFRP1, showed an expansion in proliferating cardiac cells right below the ligation i.e. in the infract zone (Figure **S5A**, **B**). When watershed areas from these two treatments were analysed carefully, these proliferating cardiac cells were rarely of arterial origin (TdTomato^+^) in PBS treated hearts (Figure **1P**). Contrastingly, watersheds of hWNT2-SFRP1 injected hearts frequently showed TdTomato^+^ EdU^+^ single artery ECs (Figure **1Q**); almost two-fold increase, compared to PBS treated hearts (Figure **1O**). Together, exogenous WNT2-SFRP1 stimulates non-regenerative MI hearts to activate artery reassembly by facilitating proliferation of pre-existing artery ECs.

### Arterial Fzd4 is essential for artery reassembly

Expansion of artery cell lineage in response to hWNT2-sFRP1 suggested arterial expression of receptors which sense surrounding Wnt ligands. Thus, we investigated the mRNA expression of *Frizzled (Fzd)* receptors in our scRNA dataset^22^ (Figure **S3A**). Among the 10 Frizzled receptors, *Fzd4* and *Fzd6* were enriched in arterial ECs (Figure **2A**). *Fzd4* showed higher expression in coronaries (as compared to Fzd6) and is known to regulate postnatal coronary artery growth^32^, making it a receptor of interest. Cryosections from P6 *Cx40CreER; Rosa26^TdTomato^* hearts were immunostained for Fzd4 to assess its colocalization with TdTomato^+^ artery ECs (Figure **2B** and Figure **S6A**). While predominantly expressed by cardiomyocytes, Fzd4 was also observed in artery ECs (Figure **2B**), prompting us to further investigate the role of arterial Fzd4 in artery reassembly.

**Figure 2:**
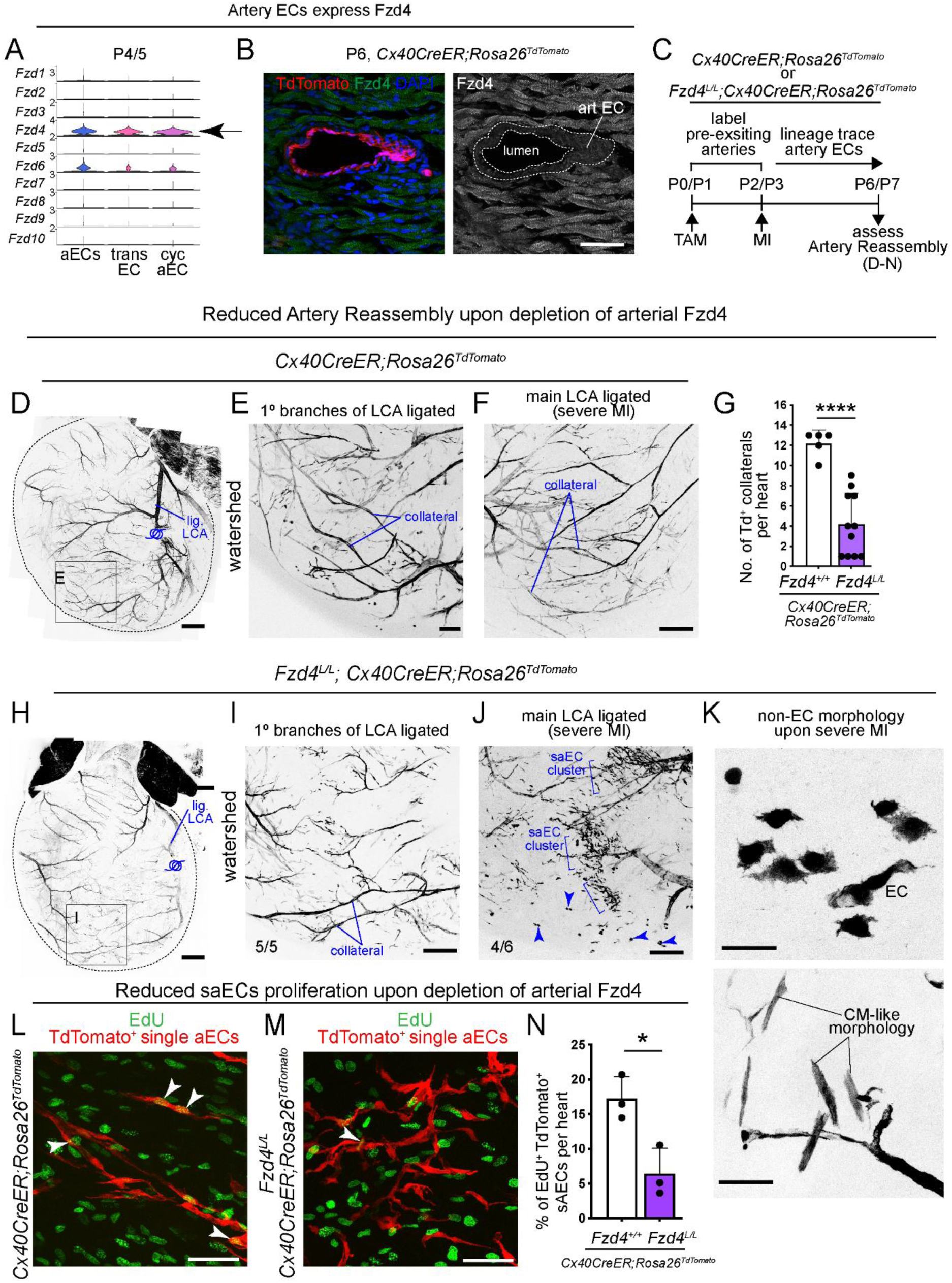
Arterial Fzd4 regulates coronary collateral formation. (**A**) Violin plot showing levels of mRNA expression of Frizzled (Fzd) receptors in neonatal hearts. (**B**) Cryosection from *Cx40CreER; Rosa26^TdTomato^* heart showing a cross-section of a coronary artery, immunostained for Fzd4 (green). TdTomato^+^ lineage traced arterial endothelial cells shown in red, nuclei labelled with DAPI in blue. (**C**) Experimental design to assess the role of arterial Fzd4 in artery reassembly. (**D**-**F**) Confocal images of neonatal *Cx40CreER; Rosa26^TdTomato^* hearts labelled and traced for pre-existing artery endothelial cells in black. **D** shows whole heart. (**E**, **F**) Images of watershed from control hearts with (**E**) moderate (inset from **D**) and (**F**) severe MI. (**G**) Quantification of TdTomato^+^ collaterals per heart. p<0.0001. (**H**-**J**) Confocal images of neonatal *Fzd4^L/L^; Cx40CreER; Rosa26^TdTomato^* hearts, labelled and traced for pre-existing artery endothelial cells (shown in black). **H** shows whole heart. (**I**, **J**) Images of watershed from control hearts with (**I**) moderate (inset from **H**) and (**J**) severe MI. Arrowheads and brackets point to single and clusters of saECs, respectively. All 5 knockout hearts undergoing moderate MI demonstrate a phenotype as shown in **I**. 4 out of 6 knockout hearts undergoing severe MI demonstrate a phenotype as shown in **J.** (**K**) Confocal image of Fzd4 depleted single artery cells, post-MI show non-EC morphology. (**L**, **M**) Confocal images of watershed regions from (**L**) control and (**M**) arterial knockouts for Fzd4, showing EdU^+^ proliferating cells in green and TdTomato^+^ lineage traced artery endothelial cells in red. Arrowheads point to EdU^+^ TdTomato^+^ proliferating single artery cells, post-MI. (**N**) Quantification of EdU^+^ proliferating single artery cells post-MI. p=0.0186. Scale bars: **B**, 50µm; **D**, **H**, 500µm; **E**, **F**, **I**, **J**, 200 µm; **K**-**M**, 50 µm. trans, transient; cyc, cycling; P, postnatal day; EC, endothelial cell; Tam, Tamoxifen; MI, myocardial infarction; lig, ligation; 1°, primary; LCA, left coronary artery; CM, cardiomyocyte; saEC, single artery endothelial cell

To assess the role of Fzd4 in collateral artery development, we lineage-labelled and simultaneously induced genetic deletion of *Fzd4* in coronary arteries with Tamoxifen at P0/P1 (Figure **2C**). Following lineage labelling, neonates were subjected to MI at P2 or P3, and hearts were harvested 4 days post-MI at P6/P7 (Figure **2C**). Hearts were processed and whole heart confocal imaging was performed at cellular resolution to reliably assess fate of lineage labelled TdTomato^+^ artery ECs in the watershed regions. Control hearts developed ∼12 TdTomato^+^ collateral arteries (Figure **2D**-**G**), while *Fzd4* knockout hearts showed 3-fold reduction in their collateral numbers (average of ∼4 collaterals) (Figure **2G**-**I**). Despite of their presence, *Fzd4* deleted single artery cells failed to build a rigorous network of collateral arteries in the watershed (Figure **2G** and **2I**). Induction of severe MI, (i.e. ligation of main LCA branch, instead of primary branches of LCA (Figure **S6B**), led to more pronounced clustering of Fzd4 depleted TdTomato^+^ single artery ECs in the collateral zone of 4 out of 6 hearts (brackets in Figure **2J** and **S6C**, **D**), and prevented formation of collateral arteries (Figure **2F**, **J** **and** **S6C**, **D**). Furthermore, *Fzd4* artery EC specific knockout watersheds often showed TdTomato^+^ cells of non-endothelial morphology (arrowheads in Figure **2J**, **2K**, Figure **S6E**-**G**), some even resembling cardiomyocytes (Figure **2K**, bottom panel and Figure **S6F** (asterisks), **S6G**), suggesting the potential role of Fzd4 in making cell fate decisions, post-injury.

### Fzd4 regulates de-differentiation of artery ECs, post-injury

Wnt signalling regulates fate of endothelial cells^25^. We checked if arterial Fzd4 is associated with de-differentiation of single artery ECs during artery reassembly. This is majorly assessed by their ability to proliferate; a feature coupled with upregulation of angiogenic/progenitor like markers^22^. Thus, we first checked the effect of arterial deletion of *Fzd4* on artery cell proliferation. This was evaluated using an EdU assay at the time of explant (at P6), 4 days post MI (Figure **2C**). Watershed areas of MI hearts were assessed for colocalization of TdTomato^+^ single artery ECs and nuclear EdU (Figure **2L**, **M**). While control hearts showed 17.2% EdU^+^ TdTomato^+^ proliferating single artery ECs, *Fzd4* knockout hearts showed a significant 2.7-fold reduction (i.e., 6.4%) (Figure **2N**). Thus, Fzd4 promotes proliferation of single artery ECs, post-MI.

Next, any changes in the expression levels of angiogenic or progenitor-like proteins such as VegfR2 or Endomucin, were evaluated in single artery ECs, post-MI (Figure **3A**). Quantifications of VegfR2 or Endomucin immunostaining of whole hearts (Figure **3B**, **C**) were performed at P5; 3 days post-MI, where typically several TdTomato^+^ saECs are observed (Figure **3A**). We first used a mouse line, *Cx40CreER; Rosa26^TdTomato^; Cx40^eGFP/+^*, which traced Cx40^+^ artery cell lineage with TdTomato and simultaneously detected Cx40 levels with eGFP (Figure **3D**). As expected^22^, single artery cells emerging from TdTomato^+^ pre-existing artery tips downregulated artery cell marker─Cx40 (arrowheads, Figure **3D**), in response to MI. As they migrated across watersheds, and along the capillaries (Figure **3E**), these single artery ECs expressed capillary marker VegfR2, at par with the surrounding capillaries (arrowheads in Figure **3F**, **G** and Figure **S7A**). In contrast, expression of VegfR2 in Fzd4 depleted single artery ECs (Figure **3H**) was significantly downregulated as compared to the neighbouring VegfR2 expressing capillary ECs (arrowheads in Figure **3I**, **J** and Figure **S7B**). Furthermore, when compared to control, expression of VegfR2 in knockout single artery ECs was significantly reduced, by 2.1-fold (Figure **3B**). A similar analysis was also performed for Endomucin expression in single artery ECs, post-MI. The emerging TdTomato^+^ single artery ECs expressed Endomucin (Figure **3K**, arrowhead in Figure **3L**, **M** and Figure **S7C**). However, *Fzd4* null single artery ECs failed to upregulate expression of Endomucin to the levels of surrounding capillaries (arrowheads in Figure **3N**, arrowheads in Figure **3O**-**P** and Figure **S7D**). Like VegfR2 (Figure **3B**), 1.8-fold reduction was observed in Endomucin expression in *Fzd4* null single artery ECs as compared to the controls (Figure **3C**). Together, our data shows that arterial Fzd4 promotes proliferation and expression of capillary/venous markers in artery ECs, during artery reassembly; two key features of arterial de-differentiation.

**Figure 3:**
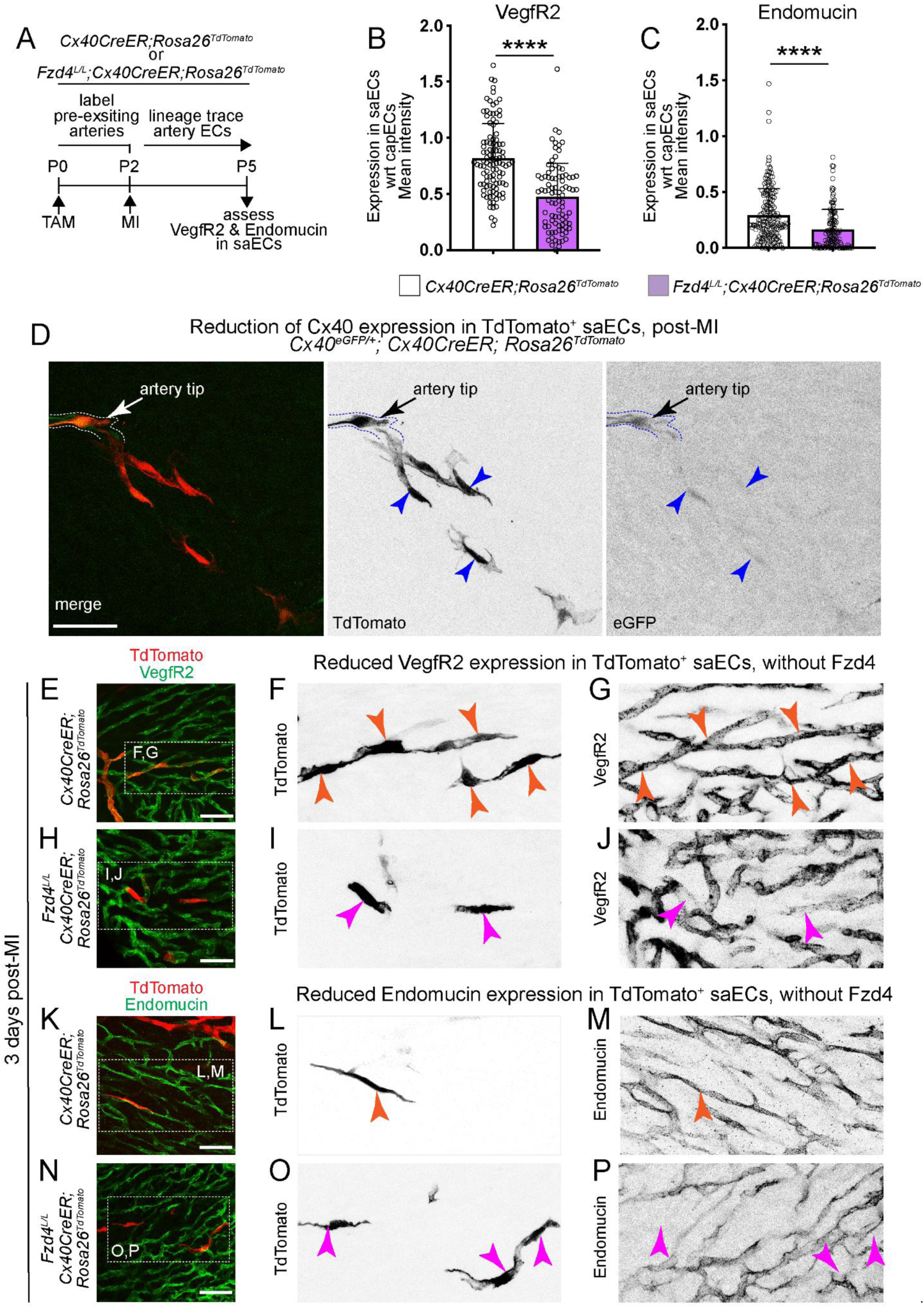
Fzd4 controls arterial de-differentiation during artery reassembly. (**A**) Experimental design to assess expression of VegfR2 and endomucin in single artery endothelial cells (saECs), post-MI. (**B**, **C**) Quantification of (**B**) VegfR2 and (**C**) Endomucin, in saECs, post-MI. p<0.0001. Each data point is a TdTomato^+^ single artery cell. N=3 and N=5 hearts were quantified for VegfR2 and Endomucin expression, respectively. (**D**) Confocal images of saECs in *Cx40CreER; Rosa26^TdTomato^; Cx40^eGFP/+^* hearts, post-MI, showing TdTomato^+^ saECs in red and expression of eGFP as a reporter for Cx40 expression in green. Arrowheads point to TdTomato^+^ saECs with low levels of eGFP signal, as compared to the artery tip. (**E**, **H**) Confocal images of post-MI watersheds, showing TdTomato^+^ saECs in red and immunostaining for VegfR2 in green in (**E**) control (*Cx40CreER; Rosa26^TdTomato^*) and (**F**) knockout (*Fzd4^L/L^; Cx40CreER; Rosa26^TdTomato^*) MI hearts. **F**, **G** are insets from **E** and show TdTomato^+^ saECs express VegfR2 (orange arrowheads). **I**, **J** are insets from **H** and show reduced expression of VegfR2 by TdTomato^+^ saECs (magenta arrowheads). (**K**, **N**) Confocal images of post-MI watersheds, showing TdTomato^+^ saECs in red and immunostaining for Endomucin in green in (**K**) control (*Cx40CreER; Rosa26^TdTomato^*) and (**N**) knockout (*Fzd4^L/L^; Cx40CreER; Rosa26^TdTomato^*) MI hearts. **L**, **M** are insets from **K** and show TdTomato^+^ saECs express Endomucin (orange arrowheads). **O**, **P** are insets from **N** and show reduced expression of Endomucin by TdTomato^+^ saECs (magenta arrowheads). Scalebars: 50µm. saECs, single artery endothelial cells; P, postnatal day; KO, knockout; cap, capillary; MI, myocardial infarction.

### Fzd4 regulates differentiation of capillaries into arteries

During embryogenesis, capillary ECs differentiate into pre-artery ECs and migrate along the coronary plexus to form intact coronary arteries^12^. This developmental process (Figure **4A**, top panel) is akin to artery reassembly^21^. We assessed the process of coronary artery development in absence of *Fzd4* in capillary ECs; the cellular source of coronary arteries during embryogenesis (Figure **4A**)^7^. *Fzd4 floxed* mice were crossed with capillary specific *ApjCreER*; *Rosa26^TdTomato^* mice. Tamoxifen induction deleted *Fzd4* from *ApjCreER*^+^ capillary ECs, and labelled them with TdTomato. Tamoxifen was administered at e10.5, before the emergence of pre-artery ECs (at ∼e12.5)^12^ and embryos were analysed at e13.5 (Figure **4B**). e13.5 hearts were immunostained for VegfR2 and CX40 to visualize capillary plexus and pre-artery/artery ECs respectively (Figure **4C**-**G**). *Fzd4* knockout embryonic hearts showed an increase in Cx40^+^ Apj^+^ artery ECs (Figure **4F**, **G**) as compared to their control counterparts (Figure **4C**, **D**). Furthermore, in contrast to controls (Figure **4D**), embryonic hearts lacking capillary *Fzd4* often demonstrated premature formation of coronary artery (Figure **4G**). 3D surface plots of confocal images (Figure **4D** for control and Figure **4G** for knockout) showed increased number of peaks for Cx40 signal, in hearts depleted for capillary specific Fzd4 (Figure **2E** and **H**). This was also evident by an overall high expression of Cx40 in the remodelling zone of knockout hearts (Figure **4I**). Expansion of coronary vascular plexus on the dorsal and ventral sides was similar between control and knockout hearts (Figure **S8**), suggesting that EC migration was unaffected in absence of capillary *Fzd4*, during coronary development.

**Figure 4:**
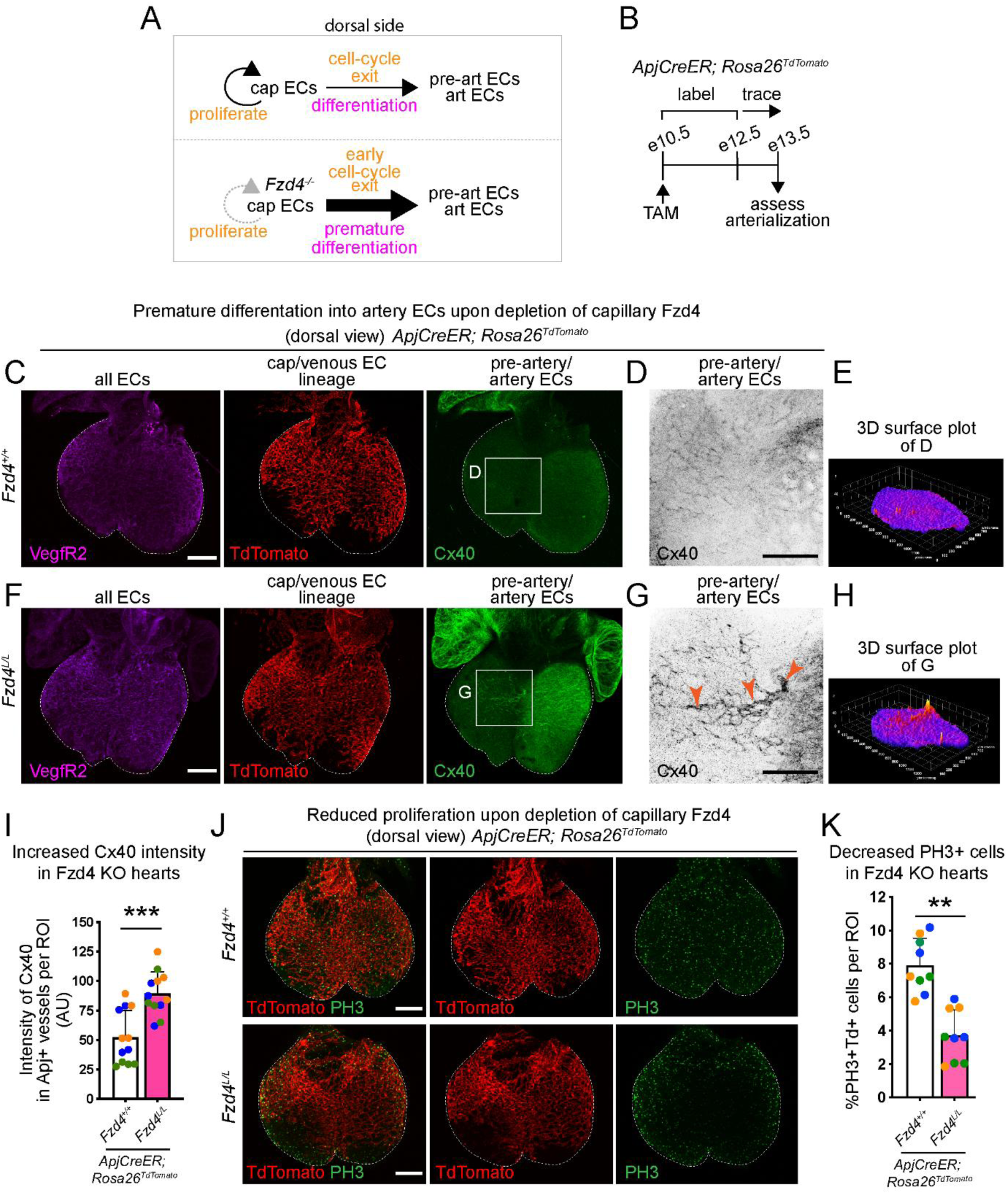
Coronary artery development is regulated by endothelial Fzd4. (**A**) Working model depicting potential role of endothelial Fzd4 in coronary artery development. (**B**) Experimental design to assess role of capillary/venous Fzd4 in coronary artery development. (**C**) Confocal image of the whole embryonic *ApjCreER; Rosa26^TdTomato^* heart immunostained for VegfR2 (in magenta), and Cx40 (in green). Apj^+^ lineage traced vessels are shown with TdTomato in red. (**D**) Inset from **C** showing immunostaining for Cx40 in black. (**E**) 3D surface plot of Cx40 signal from the image shown in **D**. (**F**) Confocal image of the whole embryonic *Fzd4^L/L^; ApjCreER; Rosa26^TdTomato^* heart immunostained for VegfR2 (in magenta), and Cx40 (in green). Apj^+^ lineage traced *Fzd4* knockout vessels are shown with TdTomato in red. (**G**) Inset from **F** showing immunostaining for Cx40 in black. (**H**) 3D surface plot of Cx40 signal from the image shown in **G**. Arrowheads point to a newly forming Cx40^+^ artery. (**I**) Quantification of intensity of Cx40 immunostaining in Apj^+^ lineage traced vessels in control and *Fzd4* knockout hearts. Each dot represents an ROI (region of interest). Each coloured dot belongs to a single heart. ROIs comprise of remodelling zone. p=0.0002. (**J**) Confocal images of embryonic *ApjCreER; Rosa26^TdTomato^* hearts with and without endothelial *Fzd4*, immunostained for PH3 shown in green. Red shows Apj^+^ vessels. (**K**) Quantification of PH3^+^ TdTomato^+^ cells. p=0.0042. Each dot is a ROIs from N=3 hearts from each genotype. ROIs comprise of remodelling zone. Scale bars: 200μm. cap, capillary; ECs, endothelial cells; e, embryonic day; PH3, Phosphohistone H3; KO, knockout

Cell cycle exit is essential for arterial fate^11,15^. To assess EC proliferation, hearts were immunostained for G2/M marker, PH3 (Figure **4J**). Lack of capillary Fzd4 led to decrease in PH3^+^ TdTomato^+^ proliferating ECs per heart, from 7.9% in control to 4.2% (Figure **4K**). Together, deletion of capillary *Fzd4* leads to an increase in Cx40^+^ cells and reduction in proliferating capillary ECs; suggesting, an early cell-cycle exit and premature differentiation into arteries (Figure **4A**, bottom panel).

To this end, data from mouse models showed that Fzd4-mediated Wnt pathway facilitates coronary artery formation during embryogenesis (Figure **S9A**) and *de novo* formation of collateral arteries, post-MI (Figure **S9B**). During development, Fzd4 maintains capillary proliferation and prevents capillary ECs from exiting cell-cycle or attain arterial fate (Figure **4** and Figure **S9A**). Similarly, upon MI, cell-autonomous Wnt activity (Figures **1**, **2**) driven by arterial Fzd4 promotes their de-differentiation (Figure **3**) in response to MI (Figures **1**-**3**, Figure **S9B**).

### Cardiovascular disease patients harbour deleterious *WLS* and *FZD4* genetic variants

Genes encoding for WNT ligands are implicated in atherosclerosis^42,43^, and acute myocardial infarction^44^. We checked if these ligands and other WNT pathway genes were associated with cardiovascular disease in South Asian Indian population. In brief, 1,177 patient exomes were analyzed, comprising two clinical categories: 335 cases of South Asian Indian hypertrophic cardiomyopathy (SAI-HCM), and 842 cases of premature coronary artery disease (SAI-pCAD). Identification of SAI-pCAD patients (with an age cut off <55 years) was primarily based on coronary angiography and the initial occurrence of myocardial infarction. Additional clinical and exome details are reported earlier and can be found here^45,46^. A robust in-house pipeline was used to analyse the whole exome sequences (Figure **S10A**). 75 WNT related genes were screened for variants (Figure **S10B**) based on cut offs for minor allele frequencies (MAFs) found in gnomAD and gnomAD SA (gnomAD South Asian) databases. These MAFs classification were based on previously published literature^47,48,49^. Accordingly, MAF <4 × 10⁻⁵ for HCM, and <1% for pCAD were considered. Interestingly, this analysis generated variants for only 4 (out of 75) Wnt pathway genes, i.e. *FZD4*, *WLS*, *WNT2 and ROR2* (Figure **S10B** and Figure **5A**). For the purposes of this study, variants of *FZD4*, *WLS* and *WNT2* were further investigated. A single *FZD4* variant (exon2: G1455T) was detected in one SAI-pCAD patient. *WNT2* variants were identified in two SAI-pCAD patients (exon5: C895T) and one SAI-HCM patient (exon5: A943G). Additionally, three *WLS* variants were detected: two variants (exon 5: G702C and exon7: T1061C) in three SAI-HCM patients, and one variant (exon3: G424A) in two SAI-pCAD patients. Thus, we identify genetic variants of *FZD4*, *WLS* and *WNT2* in South Asian Indian heart patients.

**Figure 5:**
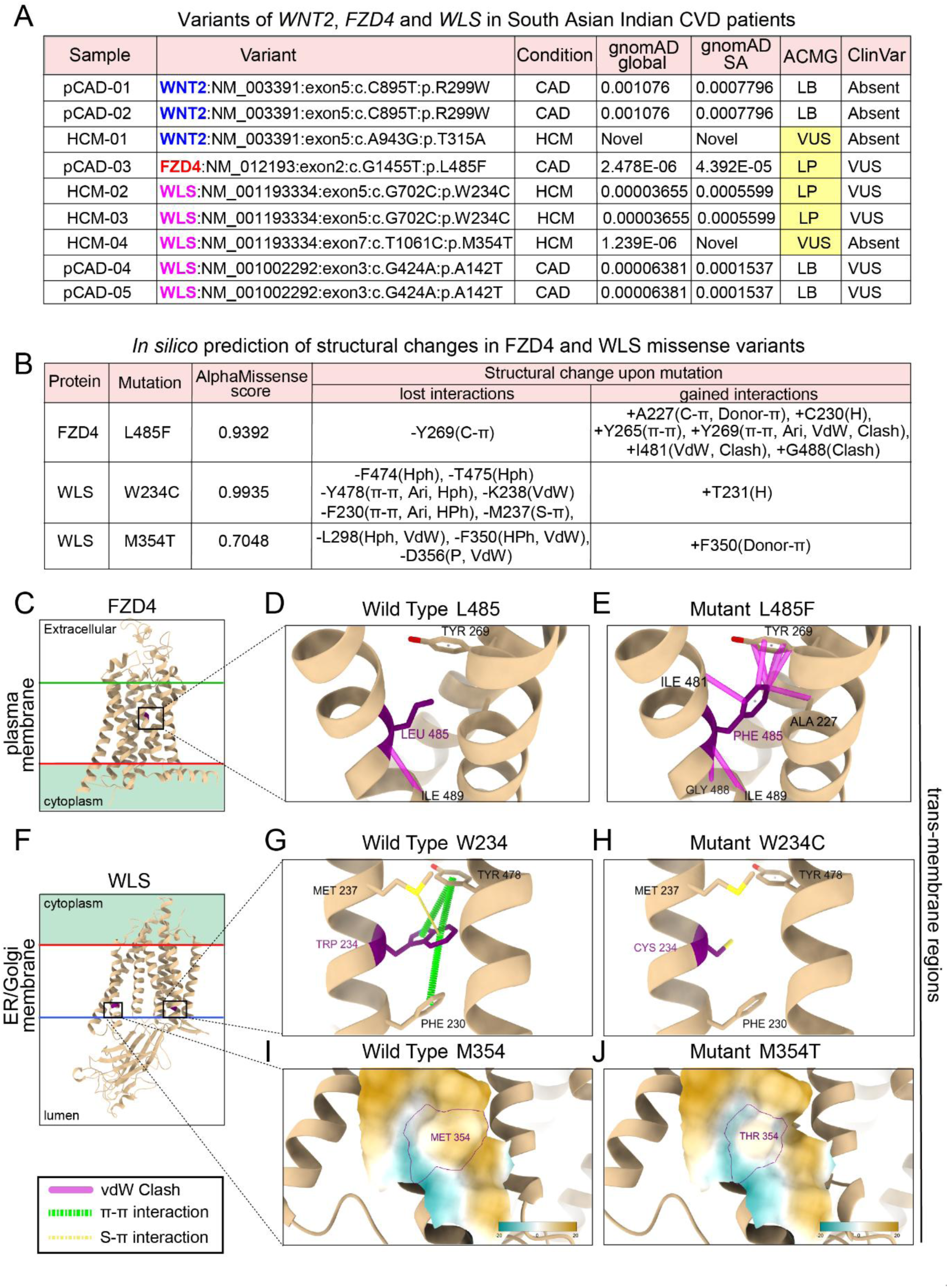
Identification of *FZD4* and *WLS* genetic variants in HCM and CAD patients. (**A**) Table listing variants of *WNT2*, *FZD4* and *WLS* genes in South Asian Indian CVD (cardiovascular disease) patients. (**B**) Table describing effects of missense mutations on gained and lost atomic interactions within FZD4 and WLS. (**C**-**J**) show predicted structural changes due to *FZD4* and *WLS* missense variants (shortlisted in **B**): (**C**-**E**) FZD4 (L485F, PDB 6BD4) and (**F**-**H**) WLS (W234C, PDB 7DRT) and (**F**, **I**, **J**) WLS (M354T, PDB 7DRT). **C** and **F** show the orientation of the proteins in the membrane, with mutated residues highlighted in purple. (**D**, **E**) Introduction of vdW clashes (magenta) in FZD4 L485F mutant. (**G**, **H**) S-π (yellow dotted line) and π-π (green dotted line) interactions present in the WLS wild-type are lost in the W234C mutant. (**I**, **J**) The molecular lipophilicity potentials in the WLS wild-type protein is reduced in M354T mutant structure. The lipophilicity potential is coloured from dark cyan (most hydrophilic) to dark goldenrod (most hydrophobic). ΔΔG value from DDmut for FZD4 L485F mutation is -1.29 (destabilizing) with a missense score of 0.9392. ΔΔG value from DDmut for WLS W234C mutation is 0.13 (stabilizing) with a missense score of 0.9935. ΔΔG value from DDmut for WLS M354T mutation is -1.14 (destabilizing) with a missense score of 0.7048. S-π, Sulphur-aromatic ring interaction; π-π, interactions between aromatic rings; vdW, van der Waals interactions; Ari, aromatic interactions; Hph, hydrophobic.

The rarity and novelty of these identified variants was assessed using gnomAD Global and gnomAD SA datasets from healthy population. Variants were classified as benign (B), likely benign (LB), pathogenic (P), likely pathogenic (LP), or variants of uncertain significance (VUS) according to ClinVar annotations and American College of Medical Genetics (ACMG) guidelines^48,50,51^ (Figure **5A**). *FZD4* (exon2: G1455T; L485F) and *WLS* (exon5: G702C; W234C) variants were classified as likely pathogenic and are previously reported in ClinVar, predominantly in individuals of South Asian ancestry. In contrast, *WLS* variant (exon7: T1061C; M354T) was absent from gnomAD SA or ClinVar and was rarely present in gnomAD Global. Based on ACMG criteria, this variant was classified as a VUS. The WNT2 variant (exon5: A943G; T315A) was classified as novel, due to its absence in gnomAD Global, gnomAD SA and ClinVar. Furthermore, this WNT2 variant was not found in South Asian Indian control datasets (i-DHANS and Indigenomes), mixed Indian and global population datasets (GenomeAsia 100K), or disease-associated datasets (Geno2MP). Thus, the shortlisted *WNT2*, *FZD4* and *WLS* variants were classified as novel, LP, and LP/VUS respectively (yellow highlight in Figure **5A**).

Missense variants result in amino acid substitutions relative to the wild-type protein. These variants often affect protein stability (proper folding) or function (including enzymatic activity, ligand binding, and protein–protein interactions), depending on the affected domain and the biochemical properties of the substituted amino acid. To evaluate the structural and functional consequences of these variants, *in silico* approaches were used to model mutant protein structures and assess the impact of each missense variant.

We analysed four missense variants—*FZD4* (L485F), *WLS* (W234C and M354T), and *WNT2* (T315A) and predicted likely pathogenicity by AlphaMissense^52^ (Figure **5A**, yellow highlight). AlphaMissense employs a structure-based deep learning framework to classify missense variants according to their pathogenic potential. Based on AlphaMissense scores, FZD4 (L485F) and WLS (W234C and M354T) were predicted to be pathogenic (Figure **5B**).

In FZD4, a membrane receptor, the L485F substitution occurs within a transmembrane helix, replacing Leucine with Phenylalanine (Figure **5C**). This introduces a bulkier aromatic side chain in place of Leucine’s Isobutyl group, leading to van der Waals clashes with neighbouring residues (A227, Y269, I481, and G488) (Figure **5B**, and compare Figure **5D**, **E**). Similarly, in WLS, primarily an endoplasmic reticulum resident transmembrane protein (Figure **5F**), substitution of Tryptophan with Cysteine at position 234 eliminates an aromatic side chain (compare Figure **5G**, **H**). Consequently, stabilizing aromatic–aromatic (π–π) stacking and sulfur–aromatic (S–π) interactions involving W234 and adjacent residues (F230, M237, and Y478) are lost (Figure **5B**). In addition, W234 is positioned near the interfacial lipid headgroup region and may normally participate in hydrogen bonding and cation–π interactions with polar lipid headgroups. These interactions are disrupted by this substitution in the mutant structure (Figure **5H**). The WLS M354T variant introduces a polar, hydrophilic Threonine in place of the hydrophobic Methionine at position 354 (Figure **5I**, **J**). Because M354 is exposed to lipid hydrocarbon tails within the transmembrane region, this substitution is predicted to disrupt membrane interactions and destabilize WLS within the lipid bilayer. Of note, the WNT2 (T315A) variant did not show major predicted effects on protein structure or stability (Figure **S10C**-**E**). Thus, 3 variants found in human WNT pathway namely, FZD4 (L485F) and WLS (W234C and M354T), were predicted to affect protein stability.

FZD4 (L485F) and WLS (W234C and M354T) are located within transmembrane regions rather than ligand-binding or protein–protein interaction domains^53,54^. Two of these three substitutions affect highly conserved residues (ConSurf scores: L485 = 8; W234 = 9) and disrupt stabilizing interatomic interactions. The WLS M354T variant specifically reduces hydrophobicity in a lipid-exposed region of the membrane. Furthermore, ΔΔG values calculated using DDMut were negative for two of these three variants, indicating reduced protein stability^55^. Collectively, these findings suggest that the identified mutations are likely to produce unstable proteins. WNT ligand secretion and signal transduction depend on WLS and FZD4 respectively. Thus, individuals harbouring L485F, W234C or M354T variants may exhibit compromised WNT signalling.

## Discussion

Using genetic lineage tracing, cell-type specific knockouts, and whole-organ imaging at single-cell resolution, we demonstrate that Wnt receptor, Fzd4 determines arterial cell fate by regulating cell-cycle progression during development and regeneration. During embryogenesis, deletion of *Fzd4* from capillaries did not affect their migration or coronary artery formation, indicating that Fzd4 specifically regulates cellular identity. Similarly, in neonatal regenerative hearts following MI, loss of arterial *Fzd4* caused artery ECs to lose their plasticity, leading to reduced proliferation and impaired de-differentiation; 2 key cellular events essential for successful execution of artery reassembly. We further showed that Wls-dependent secretion of arterial Wnt ligands is essential for their proliferation, post-MI. This confirms the cell autonomous nature of Wnt signalling in regenerative hearts. Moreover, administration of exogenous human-WNT2-sFRP1 ligand complex in non-regenerative (older) hearts was able to activate expansion of single artery EC pool via proliferation. That being said, genetic variants of *WLS* and *FZD4*, detected in human CAD patients, were predicted to be pathogenic, further underscoring the significance of WNT pathway in coronary regeneration. Together, our findings establish WNT signalling as a critical developmental pathway for coronary artery formation that is reactivated after injury to drive neo-collateral formation; a regenerative response which can be induced in non-regenerative hearts through exogenous Wnt ligand supplementation.

### Significance of endothelial de-differentiation

Arteries often rely on venous ECs for their replenishment because of their superior proliferative abilities. However, during development, venous cells switch fates to artery ECs via a *pre-artery* transient state, defined by gradual upregulation in mature artery markers and downregulation of venous as well as cell cycling genes^12^. Similarly, a recently described *pre-vein* (intermediate) state presumably determines venous identity and is marked by dual expression of arterial (Sox17) and venous (Apj) markers^56^. Co-expression of venous and arterial markers precisely indicates a de-differentiated state and offers flexibility to choose between either lineage. A molecular understanding of this transient reprogramming or EC de-differentiation can be the key to regenerative vascular biology.

Endothelial de-differentiation and proliferation are often coupled together. For instance, during development, capillary endothelial cells exit cell cycle before differentiating into arterial ECs^11,15^. Similarly, during cardiac regeneration, endothelial cells capable of de-differentiation are also proliferative. That being said, could artery de-differentiation also have roles which do not lead to proliferation? For instance, pre-artery and injured artery single cells migrate along capillaries, during development and MI respectively^7,21,57^. Both *in vitro* studies and *in vivo* analyses of vasculature show that ECs preferentially associate with cells of the same fate─ capillary ECs mix readily with capillaries, and arterial ECs with arteries^6^. This is in contrast to less favourable interactions between ECs from arteries and capillaries. Therefore, transient conversion of arterial ECs to a capillary-like state may facilitate their incorporation and migration along capillary networks. In context of our findings, this could explain clustering of Fzd4 depleted single artery cells around artery tips. It is possible that (although capable of dissociation), impaired de-differentiation of *Fzd4* deleted artery ECs, limits their ability to incorporate or migrate efficiently on capillary ECs. This incompatibility could lead to accumulation of *Fzd4* deleted artery ECs around artery tips, in response to MI. Additionally, injured *Fzd4* null artery ECs showed non-endothelial morphologies resembling mesenchymal cells. The transformed states, however temporary, could restrict a time-sensitive active contribution towards *de novo* formation of collateral arteries.

### Context-dependent endothelial Wnt signalling

The multi-faceted canonical Wnt signaling is essential for development and maintenance of vascular patterns in organisms across species^37,58^. For instance, excessive mitogenic stimulation through activation of nuclear β-catenin can drive ECs into cell-cycle arrest and promote arterial differentiation^15,59^. This could occur through recruitment of key transcriptional factors Sox17^27^ and Erg^60^, or transcription of *EphB2* and Notch ligand *Dll4*^25^; all of which promote arterial fate. Contrastingly, constitutive induction of β-catenin, genetically^26^, or through *in vitro* supplementation with canonical WNT10b ligand^35^, upregulates VegfR2, an angiogenic or capillary EC marker. Similarly, impaired endothelial β-catenin also reduces expression of VegfR2^26^, known to regulate endothelial proliferation^22^ and cell fate^11^. All these studies, together, point towards an endothelial function of canonical Wnt pathway which is heavily dependent on the developmental time, context and dosage^25,27,61^. Our data show that during coronary development, Fzd4 in capillaries keep ECs in actively cycling stage. Without Fzd4, these ECs exit cell cycle and differentiate into coronary artery ECs. This ectopic arterialization could then lead to formation of nascent artery-like structures^62^. During cardiac regeneration, arterial Fzd4 promotes capillary/venous identity and proliferation of single artery ECs, which facilitates coronary collateral formation by artery reassembly. Thus, during development of, both, coronary arteries and neonatal post-MI collaterals, endothelial Fzd4 supports an overall proliferative/de-differentiated state.

Endothelial Wnt activity is age-dependent. When subjected to MI, young ECs express higher levels of Wnt pathway molecules, as compared to older ECs^63^. Similarly, expression of Wnt ligands in isolated retinal or lung ECs decreases with age^64^. Our single-cell RNA sequencing data^22^ reveals that Wnt2 expression is enriched in arterial ECs in neonatal hearts, whereas in adult hearts, its expression shifts to a smaller capillary EC population. These age-specific Wnt activity could be because of changes in chromatin accessibility at Wnt-responsive genomic regions. Currently, we do not completely understand what regulates endothelial epigenetic landscape. It could be postnatal exposure to oxygen^65^, cell cycle states^12,59,66^, the cell-specific physical/hemodynamic microenvironment^67,68^ or a combination of all.

ECs often respond to hypoxia or ischemia through prolific expansion and angiogenesis. Upon exposure to hypoxia, retinal artery ECs do not enter cell cycle, and vascular repair is mostly driven by capillary ECs^10^. This is in contrast to neonatal hearts, where ischemia triggers de-differentiation and proliferation of pre-existing artery ECs. Interestingly, ischemia-induced coronary collateral arteries in adults originate from non-arterial sources, likely capillaries^18^. Thus, lack of blood flow (or oxygen) generates distinct EC responses. These responses could be driven by molecular heterogeneity across capillaries, veins and arteries. For instance, a crucial transcription factor of Notch pathway, RBPJ (recombination Signal Binding protein for Immunoglobulin Kappa J Region), forms a functional nuclear complex with β-catenin on RBPJ binding sites of arterial genes in artery ECs, but not in veins^69^. We also observe a canonical Wnt pathway effector Axin2, expressed in non-injured coronary artery ECs suggesting, artery ECs may require canonical Wnt signalling to maintain their arterial identity, during homeostasis^27^. Additionally, neonatal artery ECs proliferate upon WNT2 treatment. Together, these data suggest that coronary artery ECs are responsive to Wnt signalling. Currently, we do not completely understand if Wnt responsiveness relies on a particular EC sub-type or molecular heterogeneity within each sub-type. Thus, it will be interesting to know how Wnt signals from all cardiac cells converge and generate a singular effective cellular response.

### Wnt pathway genes are risk factors in CAD

We have identified 3 unique genetic variants related to Wnt pathway, which were statistically and structurally predicted to be deleterious in humans. Interestingly, these genetic variants for WNT genes, *FZD4*, *WLS* or *WNT2*, were restricted to South Asian Indian population. Previous studies^70^ have reported mutations in co-receptor, *LRP6* in Iranian^44^, Chinese^71^, and white American families^72^. These studies also indicate a possible correlation of (co-receptor) *LRP6* mutations with atherosclerosis, metabolic syndrome^44^ or hypertension^73^, all caused by elevated levels of LDL-cholesterol^74^. Similarly, genetic variations for (canonical WNT effector) *TCF7L2* polymorphisms, are positively associated with severity and mortality of CAD^75^ and is considered a cardiovascular risk factor in patients with diabetic coronary atherosclerosis^76,77^.

In atherosclerotic patients, systemic elevation of circulating WNT5A protein is additionally accompanied by increased *WNT5A* transcripts in arterial lesions^78^. A similar increase is also detected for an antagonist of WNT pathway, DKK1 (Dickkopf-1). DKK1 protein levels were increased in the serum and in advanced carotid plaques^79,80^ in patients with CAD and atherosclerosis. DKK1 sourced from platelets and ECs are thought to inflict EC activation through release of cytokines in plaque regions^79^. However, it is unclear if regional inflammation induced by DKK1 is associated with atherogenesis, plaque resolution or destabilization. The sequence of events leading to the local accumulation of WNT ligands (like WNT5A) and inhibitors (like DKK1) within atherosclerotic plaques also remain unknown. Thus, while Wnt pathway proteins upregulate upon EC dysfunction during atherosclerosis, the underlying mechanisms remain unclear.

Clinical research has so far focused on the role of Wnt genes in cellular regulation of uptake of lipid/cholesterol, which could potentially lead to inflammation^79,81^, contributing to atherosclerotic plaques. In our study we highlight a less explored role of Wnt pathway genes in endothelial cell fate decisions and their contribution to development of new arteries like coronary collaterals, upon MI, in a region away from arterial occlusion. As a proof of concept, we provided an exogeneous dose of recombinant human WNT2-sFRP1 protein complex, and rescued the vascular phenotypes arising from coronary occlusion in non-regenerative mouse hearts. Thus, Wnt-dependent endothelial fate decisions play a crucial role in coronary collateral formation and could significantly alleviate the outcome of MI-induced cardiac symptoms.

## AUTHOR CONTRIBUTIONS

SD and BB conceptualized the study. SD, PSD and SV provided resources. BB, ANS and RKZ performed *in vivo* mouse experiments. VJR performed screening for genetic variants of human *WNT* pathway genes. PSD is in charge of the work related to human CVD genomics. OG and SV performed structural analyses of human WNT pathway proteins. BB and SD wrote the manuscript. All authors contributed to editing the manuscript.

## ACKNOWLEDGEMENTS

We thank the following facilities for their technical support: Animal Care and Resource Center at NCBS, Mouse Genome Engineering Facility at NCBS, Central Imaging and Flow Cytometry Facility at NCBS.

## FUNDING

This work is supported by the following funding agencies: Department of Atomic Energy, Government of India, Project Identification No. RTI 4006 to SD and SV, Department of Biotechnology (DBT)/Wellcome Trust India Alliance Intermediate Fellowship to SD (IA/I/20/2/505205), Anusandhan National Research Foundation (ANRF, formerly SERB) Core Research Grant CRG/2023/003192 to SD, DBT grant BT/PR40323/BTIS/137/78/2023 to SV. PSD is supported by DBT (BT/PR45262/MED/12/955/2022), ANRF (CRG/2023/004193), Indian Council for Medical Research IRIS Id-2023-19831, Indo-French Centre for the Promotion of Advanced Research (IFCPAR/CEFIPRA) CSRP Project Number: 7003-1.

## Data and code availability

Scripts to analyze the effects of the missense variants in FZD4 and WLS are available at GitHub (https://github.com/isblab/wnt_signaling_mutation_analysis).

Any additional information or data will be made available upon reasonable request.

## Disclosures

None

## Methods

### Human Ethics Clearance

The studies were approved by the institutional human ethics committee (Reference number: inStem/IEC-10/001).

### Patient information

The demographics, diagnostics, inclusion/exclusion criteria, and DNA isolation protocols for exomes are detailed in our previous publication^82^.

### Single Cell RNA Sequencing analysis

Published single-cell RNA-sequencing (scRNA-seq) data from Arolkar et al., were reanalyzed in this study^22^. Raw sequencing files and Cell Ranger–processed count matrices are publicly available through the Gene Expression Omnibus (GEO) under accession number GSE210307. This dataset comprises FACS-sorted TdTomato-positive lineage-traced coronary artery endothelial cells isolated from neonatal and adult mouse hearts subjected to left coronary artery ligation or sham surgeries. Data analysis was performed in R using the same analysis pipeline and scripts provided by the original authors (https://github.com/Snehasrivatsa/scRNA). Using this pipeline, we reproduced the previously reported clustering patterns and identified cell populations via differentially expressed genes, which were consistent with those described in the original study. Using the reprocessed data, we examined the expression of Wnt ligands and Wnt receptors in lineage-traced coronary artery endothelial cells from neonatal sham and myocardial infarction (MI) hearts, as well as adult sham and MI hearts. In addition, PROGENY^83^ pathway activity analysis was performed to infer signaling pathway activity at the single-cell level. Differential pathway activity analysis was conducted to evaluate the activity of 14 curated signaling pathways across distinct arterial endothelial cell clusters in neonatal hearts.

### Experimental Model

Mouse lines used in this study were bred and maintained at the National Centre for Biological Sciences (NCBS) in accordance with Institutional Animal Care and Use Committee (IACUC) guidelines. Following mouse lines, *Cx40CreER*^84^, *ApjCreER*^85^, *Cdk1^fl/fl^* (The Jackson Laboratory, 129S(B6N)-Cdk1*^tm1Eddy^*/J, stock number 028028), *Fzd4^fl/fl^* (The Jackson Laboratory, B6;129-Fzd4*^tm2.1Nat^*/J, stock number 011078), *Wls^fl/fl^* (The Jackson Laboratory, Wls*^tm1.1Whsu^*/J, stock number 027484), *Rosa26^TdTomato^* Cre reporter line (The Jackson Laboratory, B6.Cg-Gt[ROSA]26Sortm9[CAG-TdTomato]Hze/J, stock number 007909), *Cx40^eGFP/+^* ^86^ and Axin2-d2EGFP^87^ were used in the experiments. Experiments were performed using mixed-background strains. Animals were age-matched, and both male and female mice were used in all experiments.

For postnatal experiments, a single dose of 6mg Tamoxifen (stock: 20mg/ml dissolved in corn oil) was administered to nursing mothers by intraperitoneal injection. For embryonic experiments, the same dose was administered to timed pregnant dams via oral gavage.

### Primer used for genotyping

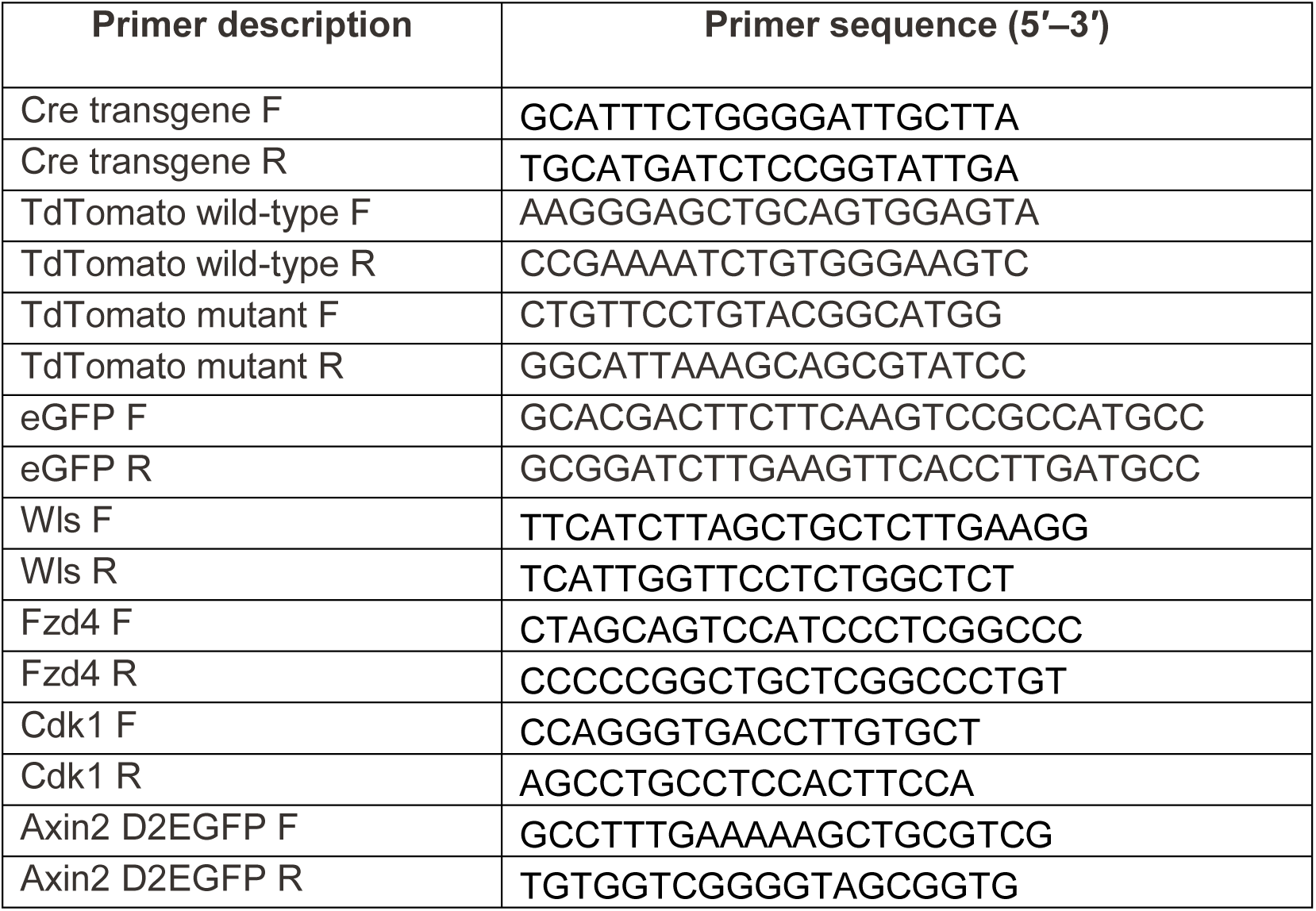

F; Forward primer sequence, R; Reverse primer sequence.

### Experimental Study Design

For all experiments, control and knockout groups were defined based on animal genotype. Only animals subjected to myocardial infarction (MI), in which either two branches of the primary left coronary artery (LCA) or the main LCA were ligated, were included in this study. Successful MI was confirmed by imaging excised hearts under a fluorescence stereomicroscope (Macro Zoom microscope MVX10; Olympus) to visualize the (ligation) knot around TdTomato positive arteries. Biological replicates from multiple litters were included in all analyses.

### Neonatal LCA ligation

Surgical procedures were modified from Mahmoud et al.^88^ and performed under sterile conditions. Neonatal mice (P2/P3 or P7) were wrapped in precooled gauze and placed on ice for 4 minutes to induce hypothermic cardiac arrest. Loss of consciousness was confirmed by toe-pinch reflex prior to surgery. Mice were positioned supine on a cold ice pack, wrapped with a sterile surgical drape. The mice were prepared using betadine followed by 70% ethanol. A left anterolateral thoracotomy was performed under a dissecting microscope by making an incision at the fourth intercostal space. The heart was visualized, and the left coronary artery (LCA) was identified and ligated using an 8-0 nonabsorbable Prolene suture (catalog no. NW3322; Ethicon Ethilon) attached to a needle. Following ligation, the ribs, intercostal muscles, and skin were closed separately using the same 8-0 Prolene suture. After surgery, neonates were transferred to a 37°C warm plate until full recovery and return of spontaneous movements. Recovered pups were returned to the parent cage and monitored closely over the subsequent days.

### Intramyocardial injection of hWNT2-sFRP1

Intramyocardial injection of hWNT2-sFRP1 was performed while performing LCA ligation surgery on neonatal P7 mice. Following ligation, a single dose of hWNT2-sFRP1 was applied, 15 μL (55ng) of hWNT2-sFRP1 (R&D System; Cat:1117-WN-0101) diluted in 1X phosphate buffered saline or vehicle only (1X PBS) was injected intramyocardially into the watershed region.

### Whole mount immunostaining of neonatal hearts

Neonatal whole hearts were dissected, washed in 1X PBS, and immediately fixed in 4% paraformaldehyde at 4°C for 1 hour with gentle rocking. Hearts were then washed three times for 15 mins each in 1X PBS at 4°C. Hearts were permeabilized over 3–4 days by washing in 0.5% PBT (1X PBS containing 0.5% Triton X-100) using 20 volumes per wash, 5–6 washes, with buffer changes every 2 hours at 4°C. Primary antibody staining was performed by incubating hearts in the appropriate primary antibody dilutions prepared in at least 5 volumes of 0.5% PBT, overnight at 4°C with gentle shaking. Hearts were then washed in 20 volumes of 0.5% PBT for 12 hours (5–6 washes) at 4°C. Secondary antibody staining was carried out by incubating hearts in a 1:250 dilution of secondary antibodies prepared in 0.5% PBT, with gentle shaking. This was followed by extensive washing in 20 volumes of 0.5% PBT for 4 days, with buffer changes every 2 hours at 4°C. All washes were performed with continuous gentle agitation. Prior to clearing, hearts were washed 3–4 times in 1X PBS. Hearts were then cleared in at least 2 volumes of antifade mounting medium (https://www.jacksonimmuno.com/technical/products/protocols/anti-fade) for 2 hours, protected from light, at room temperature. Whole hearts were stored at −20°C until imaging. Post-staining washes were performed in 15- or 50-mL tubes, whereas antibody and mounting incubations were carried out in 1.5-mL microcentrifuge tubes.

For EdU labeling, neonatal mice were injected with 10 µl per gram body weight of 10 mM 5-ethynyl-2′-deoxyuridine (EdU) prepared in culture grade 1X PBS, 1 hour prior to collection of hearts. EdU detection was performed following secondary antibody staining using the Click-iT EdU Imaging Kit (catalog no. C10340; Invitrogen) according to the manufacturer’s instructions.

### Whole mount immunostaining of embryonic hearts

Embryos were dissected from pregnant dams, washed in 1X PBS, and immediately fixed in 4% paraformaldehyde at 4°C for 1 hour with gentle rocking. Embryos were then washed three times for 15 minutes each in 1X PBS at 4°C. Hearts were dissected from embryos and permeabilized for 1 day by washing in 0.5% PBT (1X PBS containing 0.5% Triton X-100) using 20 volumes per wash, 5–6 washes, with buffer changes every 2 hours at 4°C. Primary antibody staining was performed by incubating hearts in the appropriate primary antibody dilutions prepared in at least 5 volumes of 0.5% PBT, overnight at 4°C with gentle shaking. Hearts were then washed in 20 volumes of 0.5% PBT for 12 hours (5–6 washes) at 4°C. Secondary antibody staining was carried out by incubating hearts in a 1:250 dilution of secondary antibodies prepared in 0.5% PBT, with gentle shaking, followed by the same washing scheme described above. All washes were performed with continuous gentle agitation. Finally, hearts were transferred to antifade mounting medium (https://www.jacksonimmunumber.com/technical/products/protocols/anti-fade) and oriented with the dorsal or ventral side in a glass chamber slide, ready for confocal imaging. Post-staining washes were performed in 15-mL tubes, whereas antibody incubations were carried out in 1.5-mL microcentrifuge tubes.

### Confocal imaging of neonatal hearts

Lineage-traced or immunostained neonatal whole hearts were mounted on a double-concave microscope slide (catalog no. 7104; Sail Brand) with a coverslip (thickness 0.13–0.17 mm; catalog no. 48367-059; Thermo Scientific) such that the ventral watershed region faced upward. Whole hearts were imaged using 4X, 10X, or 60X oil-immersion objectives with an Olympus FV3000 upright confocal microscope. Image acquisition was performed using FV31S-SW software to collect multiple z-stacks. Image processing was carried out in Fiji/ImageJ, and individual fields were stitched to generate whole-heart images using Adobe Photoshop.

### Immunofluorescence on cryosections

Neonatal whole hearts were dissected, washed in 1X PBS, and immediately fixed in 4% paraformaldehyde at 4 °C for 1 hour with gentle rocking. Hearts were then washed three times for 15 minutes each in 1X PBS at 4°C and subsequently incubated in 30% sucrose at 4°C for 3–5 days for cryoprotection. Hearts were removed from sucrose, embedded in OCT compound (Optical Cutting Temperature; Fisher Healthcare, catalog no. 4585), snap-frozen on dry ice, and stored at −20 °C for 24 h before transfer to −80 °C. Cryosections (10 µm) were prepared and stored at −80 °C until further use. For immunostaining, cryosections were thawed at room temperature for 15 minutes, followed by two washes in 0.5% PBT. Sections were incubated with primary antibodies diluted in 0.5% PBT overnight at 4 °C. Slides were then washed in 0.5% PBT and incubated with secondary antibodies diluted in 0.5% PBT for 3 hours at room temperature, followed by additional washes. Sections were mounted with coverslips, using Vectashield, sealed with nail polish, and were imaged.

### Secretome analysis

This protocol was adapted from Barallobre-Barreiro et al.^41^ and employs a sequential extraction strategy using three buffers to isolate distinct protein pools from tissue. The buffers used were: NaCl buffer (0.5 M NaCl, 10 mM Tris-HCl, 25 mM EDTA), SDS buffer (0.1% SDS, 25 mM EDTA), and GuHCl buffer (4 M guanidine hydrochloride, 50 mM sodium acetate, 25 mM EDTA), all prepared in molecular grade water. Sequential incubation of tissue in these buffers enables extraction of different protein fractions. The NaCl buffer extracts loosely bound and soluble extracellular matrix (ECM) proteins as well as actively secreted proteins due to its low ionic strength. The SDS buffer solubilizes membrane-associated and intracellular proteins without disrupting ECM proteins at the concentration used. The GuHCl buffer extracts tightly bound and heavily cross-linked ECM proteins.

Neonatal whole hearts were dissected, weighed, diced into 3–4 pieces, and transferred to 1.5-mL microcentrifuge tubes. Approximately 10mg of tissue was sufficient for detection of Wnt proteins by western blotting. Samples were washed five times with 500µL ice-cold 1X PBS. PBS was pre-chilled, and a protease inhibitor cocktail (1:100) was freshly added to all buffers immediately before use. NaCl buffer was added at 10X volume-to-weight (V/W) relative to tissue mass, and samples were incubated at 25°C for 4 hours with shaking at 200 rpm, ensuring that the temperature remained between 20–25°C throughout the incubation. Extracts were transferred to fresh tubes and centrifuged at 3,000g for 10 minutes at 4°C, then stored at −20°C until further use. Because Wnt proteins are sensitive to high centrifugal forces, centrifugation speeds were maintained at 3,000g. However, if visible debris remained, the speed was increased to 16,000 g to remove particulate matter. The remaining pellet was gently washed with 100–200 µL NaCl buffer, and the wash buffer was discarded without disturbing the tissue. SDS buffer was then added to the pellet at 10X V/W, and samples were incubated at 25°C for 2 days with shaking at 200 rpm. Vigorous pipetting was avoided to prevent foaming. Extracts were collected, centrifuged at 3,000g for 10 minutes at 4°C, and stored at −20°C. The residual pellet was washed with 100–200 µL distilled water to ensure complete buffer removal. GuHCl buffer was subsequently added at 5X V/W, and samples were incubated at 25°C for 4 days with shaking at 600 rpm. Extracts were transferred to fresh tubes, centrifuged at 3,000g for 10 minutes at 4°C, and stored at −20°C until analysis.

Protein concentration was determined using the BCA Protein Assay Kit, using 10µL of each sample. For GuHCl-containing samples, 5µL of extract was mixed with 5µL RIPA buffer to reach a final volume of 10µL, as high concentrations of GuHCl are incompatible with SDS–PAGE. After addition of sample loading buffer, lysates were centrifuged for a few seconds, and not for extended periods. Samples were heated at 70°C for 10 minutes prior to electrophoresis. Proteins were separated by SDS–polyacrylamide gel electrophoresis and transferred to nitrocellulose membranes at 120 V and ∼300 mA for 1 hour 30 minutes to 1 hour 45 minutes. Transfer efficiency was confirmed by Ponceau S staining. Membranes were then incubated with primary antibodies Wnt2 (1:4000 diluted in 5% blotto) and β-actin (1:2000 diluted in 5% blotto), washes were done with 1XPBS in 0.1% Tween-20, followed by detection with horseradish peroxidase–conjugated secondary antibodies.

### Antibodies

Primary antibodies: Anti-Cx40 (1:300, catalog number CX40-A, Alpha Diagnostics Int, Inc), Anti-VegfR2 (1:200, catalog number AF644, R&D Systems), Anti-Endomucin (1:300, catalog number 14-5851-82, Invitrogen), Anti-Fzd4 (1:300, catalog number MAB 194-050), Anti-β-catenin (1:200, catalog number D10A8, CST), Anti-PH3 (1:300, catalog number 06-570, Sigma), Anti-GFP (1:300, catalog number ab13970, abcam).

Secondary antibodies: Alexa fluor-conjugated antibodies (488, 555, and 633) from Invitrogen were used at 1:250 dilutions.

### Quantification

All statistical assessments were done using unpaired t-tests. All p-values are included in the figure legends.

#### Collateral number

Collateral number was quantified at 10X magnification using an Olympus FV3000 upright microscope. TdTomato (lineage traced) collaterals were counted at the time of imaging by scanning through z-axis within watershed region of whole-heart. A continuous TdTomato positive vessel extending from the ligated artery and connecting to the RCA or septal tree was counted as one collateral.

#### saECs number and proliferation

saECs (TdTomato⁺ and EdU⁺TdTomato⁺) were quantified from images acquired using a 60X oil-immersion objective. Multiple fields of views (FOVs) were collected from each heart. A FOV was included for quantification only if at least three saECs were present. Image slices used for quantification were matched and comparable between experimental groups. In regenerative hearts, a total of 1,249 TdTomato⁺ saECs were counted in control hearts, 1,328 saECs in *Wls* knockout hearts, 1,064 sAECs in *Fzd4* knockout hearts, and 348 saECs in *Cdk1* knockout hearts. In non-regenerative hearts, 381 saECs were quantified from 1X PBS injected hearts and 488 saECs from hWNT2-sFRP1 treated hearts.

#### Intensity of VegfR2 and Endomucin in saECs

VegfR2 and Endomucin staining in saECs were quantified from images acquired using 60X oil-immersion objective with a z-step size of 1.90 µm. Using Fiji-ImageJ, a region of interest (ROI) was manually drawn to mark the boundary of individual saECs. Only saECs with the majority of their volume contained within a single optical slice were included for analysis. Mean fluorescence intensity for VegfR2 and Endomucin was measured by applying the same ROI to the corresponding fluorescence channels. Within the same optical plane, fluorescence intensity from three randomly selected capillary segments was measured and averaged to serve as an internal control. Background fluorescence was subtracted from both saECs and capillary measurements. Mean fluorescence intensity of VegfR2 and Endomucin was then normalized to capillary intensity within the same plane for saECs of both control and *Fzd4* knockout hearts. For VegfR2 analysis, 105 saECs from control hearts and 88 saECs from *Fzd4* knockout hearts were quantified. For Endomucin analysis, 184 saECs from control hearts and 127 saECs from *Fzd4* knockout were quantified.

#### LCA width

LCA width for the main and primary branches were measured using a line selection tool on Fiji-ImageJ. Quantification was done from 10X confocal images, for main branch (where visible) and primary branches of the LCA in all hearts and genotypes by averaging 3 separate measurements along the length of the branch. The analysis was done for 5 control, 4 *Cdk1* knockout, 3 *Fzd4* knockout and 2 *Wls* knockout non-injured hearts.

#### Cx40 intensity in embryonic hearts

Cx40 fluorescence intensity was quantified from fields of views (FOV) imaged using a 10X objective with 2X digital zoom on an Olympus inverted confocal microscope. All hearts were mounted in the same orientation, and four FOVs per heart were acquired from the dorsal side of the left ventricle with a z-step size of 5 µm. Three consecutive optical slices were used for quantification. TdTomato fluorescence channel was thresholded using the Otsu method, converted to a binary mask, and subjected to morphological closing. This mask was used as a selection to measure mean fluorescence intensity in the Cx40 channel. Background fluorescence was subtracted from the measured intensities. Image slices used for quantification were matched and comparable between experimental groups. For each FOV, Cx40 mean fluorescence intensity was multiplied by the corresponding mask area to obtain integrated Cx40 signal, which was then summed across slices and normalized to the total TdTomato positive vessel area. The resulting Cx40 fluorescence intensity per TdTomato vessel pixel was compared between control and knockout hearts.

### Proliferation in *ApjCreER; Rosa26^TdTomato^* embryonic hearts

Capillary proliferation was quantified from fields of view (FOV) imaged using a 10× objective with 2× digital zoom on an Olympus inverted confocal microscope. For each FOV, four 200 µm × 200 µm regions of interest (ROIs) were used for quantification across image stacks acquired with a z-step size of 5 µm. PH3⁺ApjCreER; TdTomato⁺ cells and total PH3⁺ cells were counted within each ROI per FOV. In embryonic hearts, a total of 1428 PH3⁺ cells were quantified in control hearts and 1557 PH3⁺ cells in Fzd4 knockout hearts.

### Vessel coverage in *ApjCreER; Rosa26^TdTomato^* embryonic hearts

TdTomato coverage was quantified from field of views (FOVs) imaged using a 4X objective with 1.5X digital zoom on an Olympus inverted confocal microscope. All hearts were mounted in the same orientation. The VegfR2 fluorescence channel was used to delineate the heart boundary and served as the region of interest (ROI). The TdTomato fluorescence channel was thresholded using the Otsu method, and area fraction was calculated within the whole-heart ROI. Area fraction was used as a measure of TdTomato coverage. TdTomato coverage was compared between control and *Fzd4* knockout embryonic hearts on both the dorsal and ventral sides.

### *In silico* prediction of structural changes upon mutation in FZD4, WLS and WNT2

The PDBe (Protein Data bank in Europe) REST (Representational State Transfer) API (Application Programming interface) was used to find the most complete and highest resolution available experimental structures for FZD4 (PDB 6BD4) and WLS (PDB 7DRT)^89,53,54^. WNT2 structure was obtained from the AlphaFold database^52,90^. AlphaMissense scores of mutations (0-0.34; LB, 0.34-0.564; Uncertain and 0.564-1.00; LP) were obtained from the AlphaFold Protein Structure Database^52,90^. DDMut was used to predict the mutant structure via their API^55^. ChimeraX; visualization tool; was used to add hydrogen atoms to the structures of both wild-type and mutant proteins, followed by Arpeggio to enumerate their inter-atomic interactions^91,92^. The orientations of the proteins in the membrane shown in the figure were obtained from the Orientations of Proteins in Membranes database^93^. ConSurf (1-3; variable, 4-6; intermediate and 7-9; highly conserved) was used to generate amino acid conservation scores for both FZD4 and WLS^94^.

### Whole-exome sequencing and variant analysis in human

Genomic DNA was isolated from peripheral blood mononuclear cells (PBMCs) obtained from patients. Exome libraries, comprising coding exons and selected untranslated regions (UTRs), were prepared from genomic DNA using the SureSelect V5 Enrichment Kit (Agilent). Next-generation sequencing (NGS) was performed to generate 100bp paired-end reads with an average sequencing depth of 100X using a previously published pipeline^82^.

Sequenced data were processed using an in-house exome analysis pipeline^82^. Raw sequencing reads were quality-trimmed to remove low-quality bases and adapters. The processed FASTQ files were aligned to the human reference genome (GRCh38) using the Burrows–Wheeler Aligner (BWA-MEM, version 0.7.17). The resulting SAM files were converted to BAM format, and PCR duplicates were marked and removed using Picard tools (version 2.9.0). Variant calling of South Asian Indian (SAI) premature coronary artery disease (SAI-pCAD) was performed using Mutect2 from the Genome Analysis Toolkit (GATK version 4.1.9.0), with a minimum read depth of 3 and a variant allele frequency (VAF) threshold of 2%. In contrast, variant calling of SAI hypertrophic cardiomyopathy (SAI-HCM) was performed using Haplotype Caller from GATK version 4.1.9.0. Identified variants were annotated for gene context, functional consequence, and population allele frequency using ANNOVAR^95^, with reference to the gnomAD database. Variants were further filtered based on predicted pathogenicity using *in silico* tools, including Sorting Intolerant From Tolerant (SIFT), which assesses evolutionary conservation; Polymorphism Phenotyping v2 (PolyPhen-2), which predicts structural and functional impact; and Combined Annotation Dependent Depletion (CADD) scores, with a Phred score >20 indicating variants within the top 1% of predicted deleterious changes. Rare and novel variants were prioritized for downstream analysis based on minor allele frequency (MAF) thresholds: <4 × 10⁻⁵ for HCM, and <1% for pCAD, according to gnomAD v4.1^47,48,49^. Finally, variants were classified as benign (B), likely benign (LB), pathogenic (P), likely pathogenic (LP), or variants of uncertain significance (VUS) based on ClinVar annotations and American College of Medical Genetics and Genomics (ACMG) guidelines (4, 5). Variants with conflicting classifications between ClinVar and ACMG (P/LP vs VUS) were assigned as VUS^48, 51^.

## Supplemental figures and legends

**Supplementary Figure 1:**
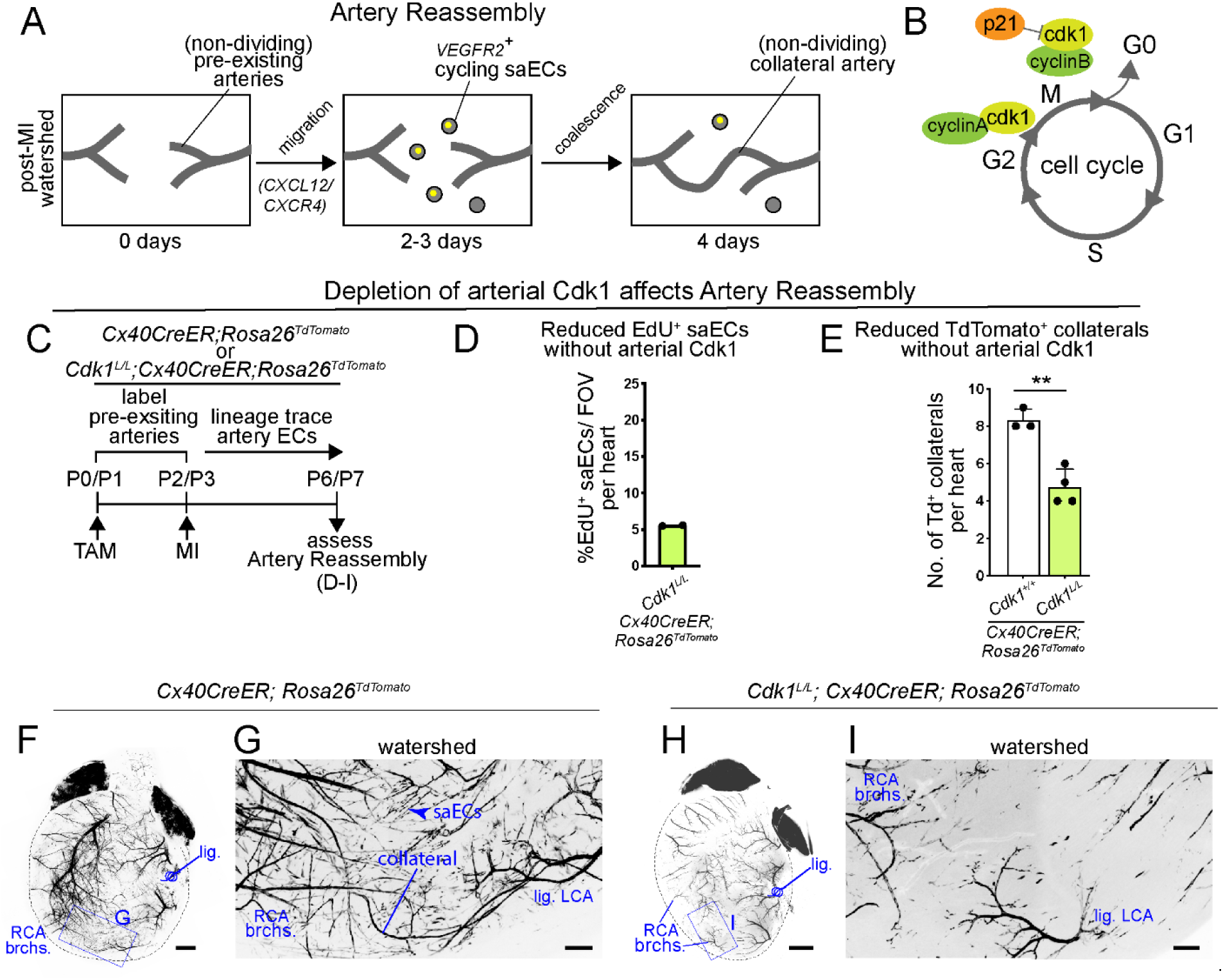
Role of arterial Cdk1 in artery reassembly. (**A**) Schematic showing cellular events associated with artery reassembly. (**B**) Schematic showing Cdk1 in mammalian cell cycle. (**C**) Experimental design to assess the role of arterial Cdk1 in artery reassembly. (**D**) Quantification of EdU^+^ proliferating single artery cells post-MI. (**E**) Quantification of TdTomato^+^ collaterals per heart. p=0.0018 (**F**) Confocal image of *Cx40CreER; Rosa26^TdTomato^* whole heart with arteries in black. **G** is an inset from **F** and shows the watershed region. (**H**) Confocal image of *Cdk1^L/L^; Cx40CreER; Rosa26^TdTomato^* whole heart with arteries in black. **I** is an inset from **H** and shows the watershed region. Scale bars: **F**, **H**, 500µm; **G**, **I**, 200µm. TAM, Tamoxifen; P, postnatal day; MI, myocardial infarction; Td, TdTomato; saEC, single artery endothelial cells; RCA, right coronary artery; brchs., branches; lig, ligation; LCA, left coronary artery; FOV, field of view

**Supplementary Figure 2:**
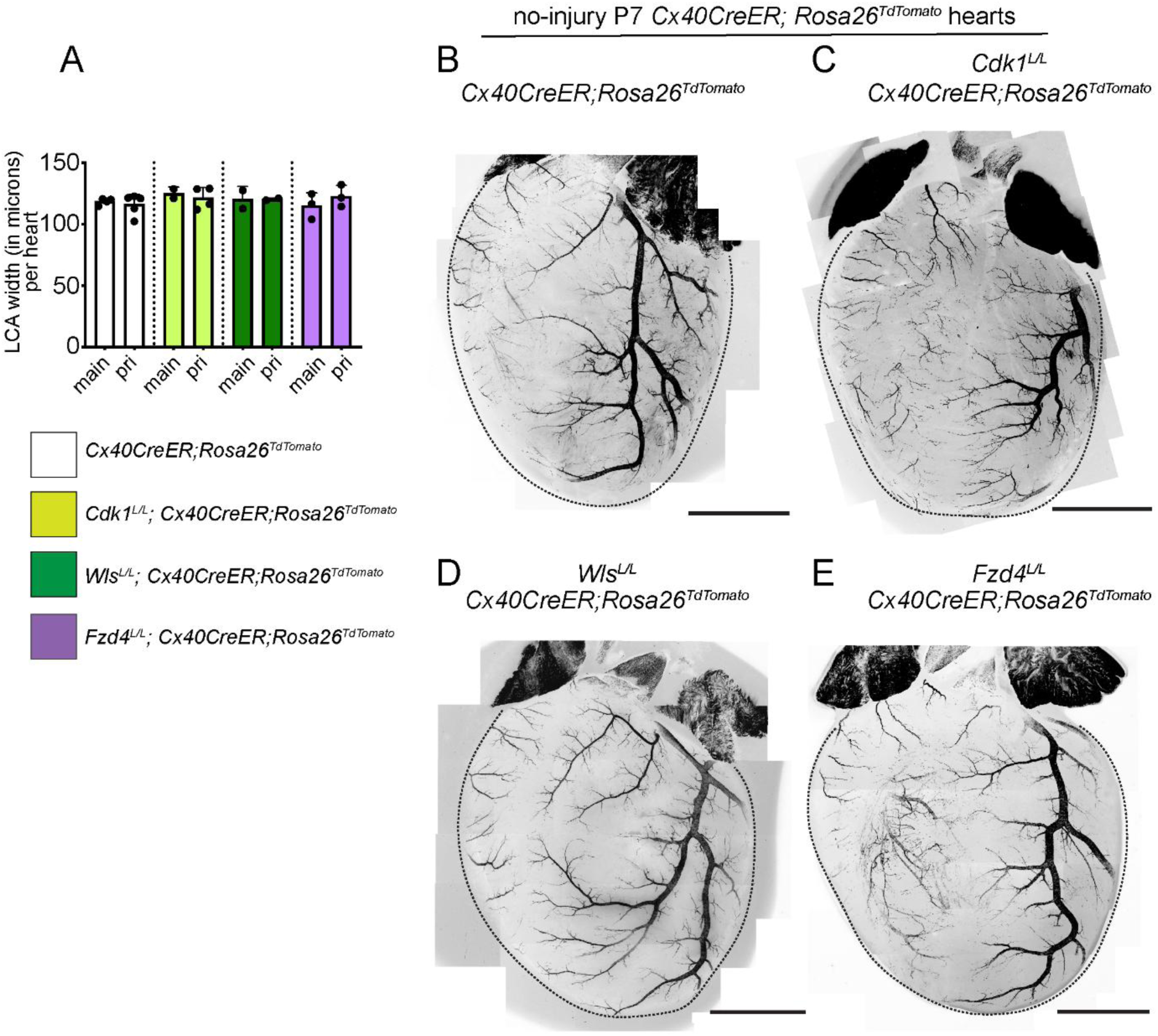
Coronary artery network of uninjured hearts, depleted in proliferation-driving genes. (**A**) Quantification of LCA widths of different genotypes. Each data point is the average value from a single heart. p values: CT versus *Cdk1* KO, p values (main) 0.3173 and (primary branch) 0.3848; CT versus *Wls* KO, p values (main) 0.8620 and (primary branch) 0.3414; CT versus *Fzd4* KO, p values (main) 0.6351 and (primary branch) 0.3555. (**B**-**E**) Confocal images of whole hearts from (**B**) wild-type or mice depleted for arterial (**C**) *Cdk1* (**D**) *Wls* and (**E**) *Fzd4*. Tamoxifen was injected at P0/1, hearts were analysed at P6/7. LCA, left coronary artery; pri, primary; KO, knockout; P, postna tal day. Scale bar: 1.5mm

**Supplementary Figure 3:**
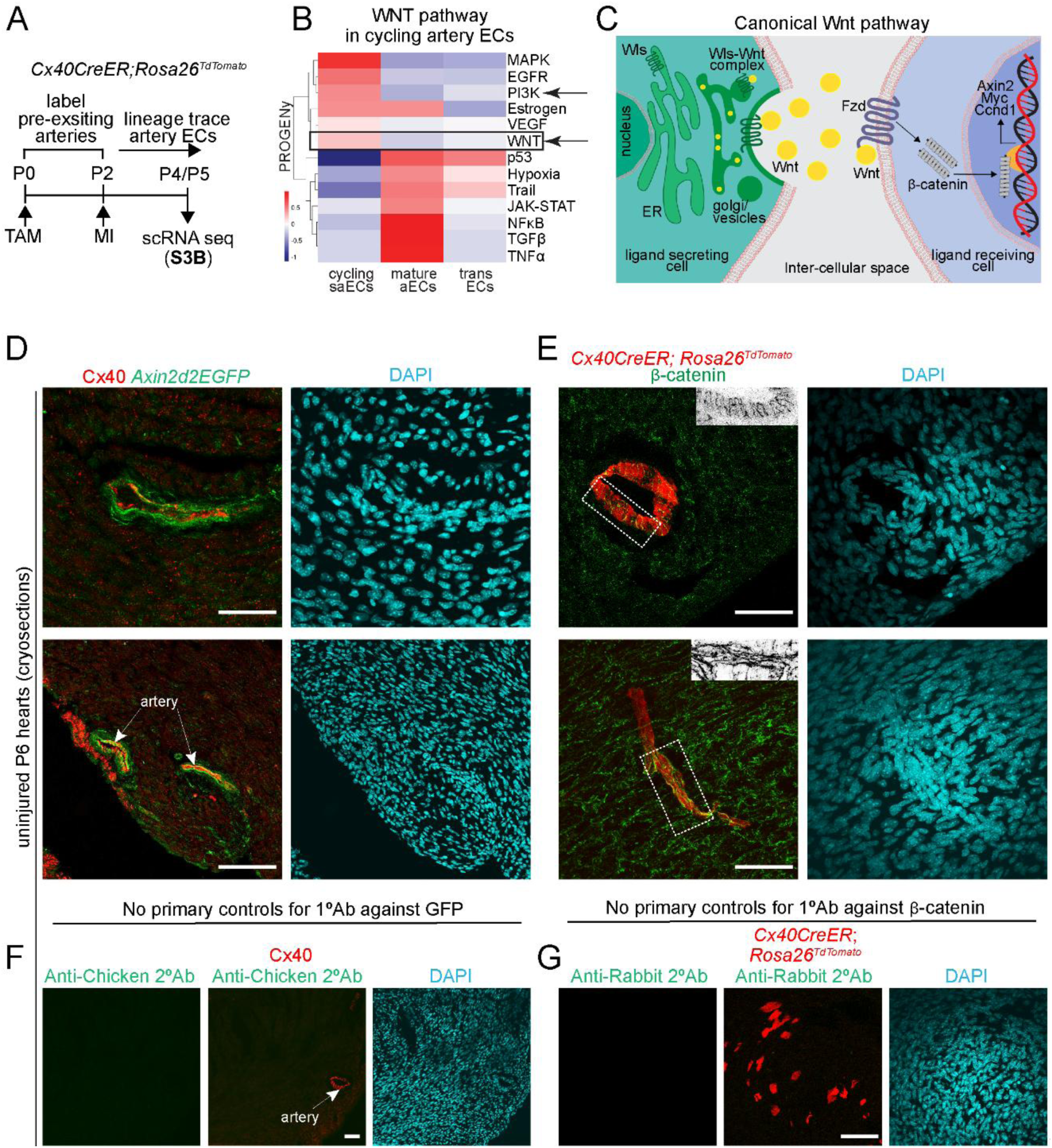
Wnt activity in coronaries. (**A**) Experimental design for single cell RNA sequencing (scRNA seq) data shown in **B**. (**B**) PROGENy analyses of coronary artery endothelial cells. (**C**) Schematic showing canonical Wnt pathway. (**D**) Cryosections from *Axin2d2EGFP* hearts showing expression of GFP reporter (green) in Cx40^+^ coronaries (red). DAPI labels nuclei in blue. (**E**) Cryosections from *Cx40CreER; Rosa26^TdTomato^* hearts showing expression of β-catenin (green) in TdTomato^+^ coronaries (red). DAPI labels nuclei in blue. Insets (marked as dotted rectangles) at the top right corner show β-catenin signal in grey scale. (**F**) Secondary alone control for immunostaining shown in **D**. (**G**) Secondary alone control for immunostaining shown in **E**. Scale bars: 50µm. TAM, Tamoxifen; MI, Myocardial infarction; trans, transient; aEC, artery endothelial cell; saEC, single artery endothelial cell; 1°, primary; 2°, secondary; Ab, antibody.

**Supplementary Figure 4:**
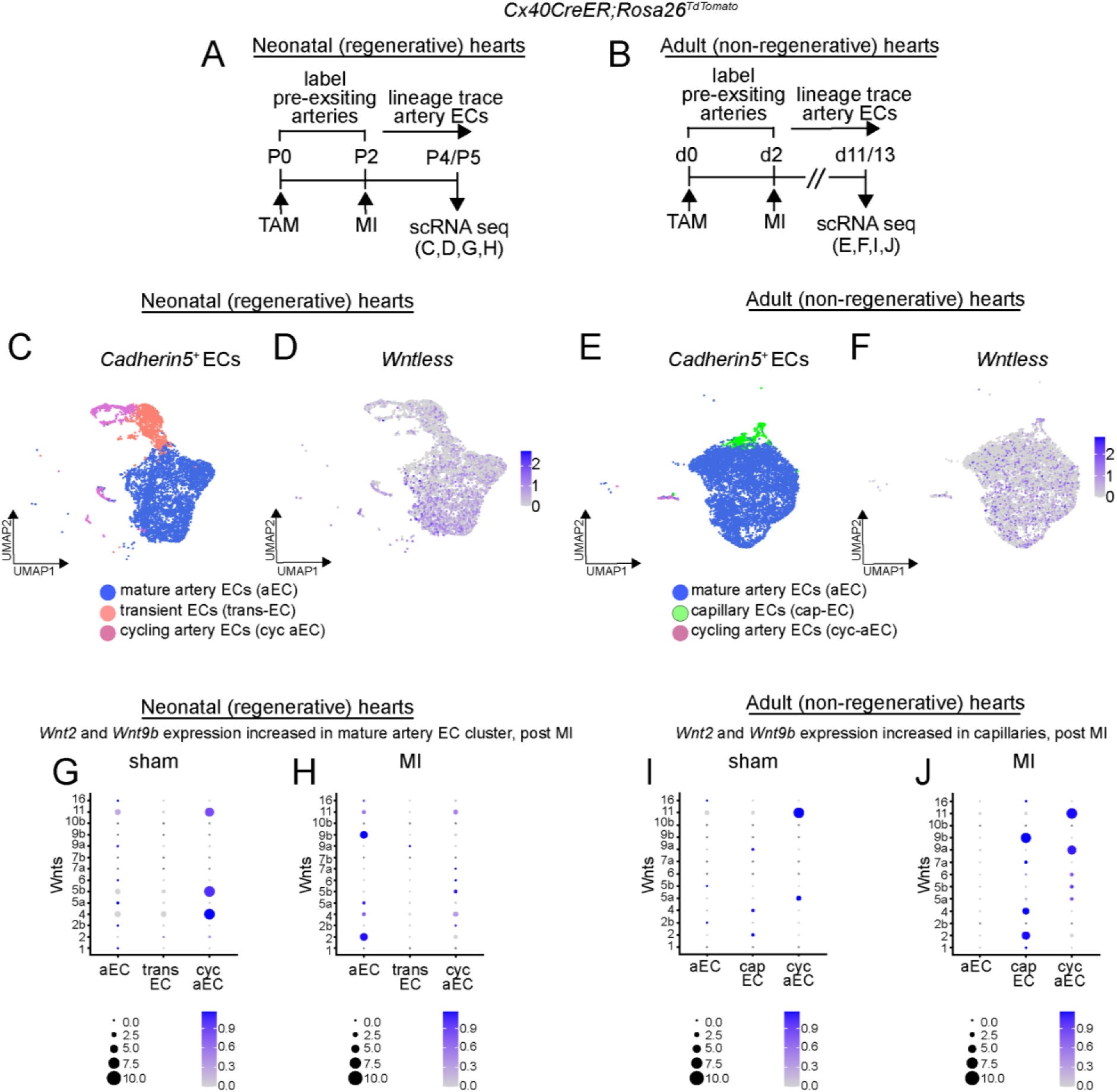
Age-dependent differential expression of Wnt pathway genes. (**A**) Experimental design for single cell RNA sequencing (scRNA seq) data shown in **C**, **D**, **G**, **H**^22^. (**B**) Experimental design for scRNA seq data shown in **E**, **F**, **I**, **J**^22^. **C** shows *Cadherin5*^+^ neonatal coronary endothelial cell clusters. (**D**) Feature plot representing *Wls* expression in all *Cadherin5*^+^ neonatal ECs. **E** shows *Cadherin5*^+^ adult coronary endothelial cell clusters. (**F**) Feature plot representing *Wls* expression in all *Cadherin5*^+^ adult ECs. (**G**, **H**) Dot plots showing expression of *Wnt* ligands in neonatal (**G**) sham and (**H**) MI hearts. (**I**, **J**) Dot plots showing expression of *Wnt* ligands in adult (**I**) sham and (**J**) MI hearts.

**Supplementary Figure 5:**
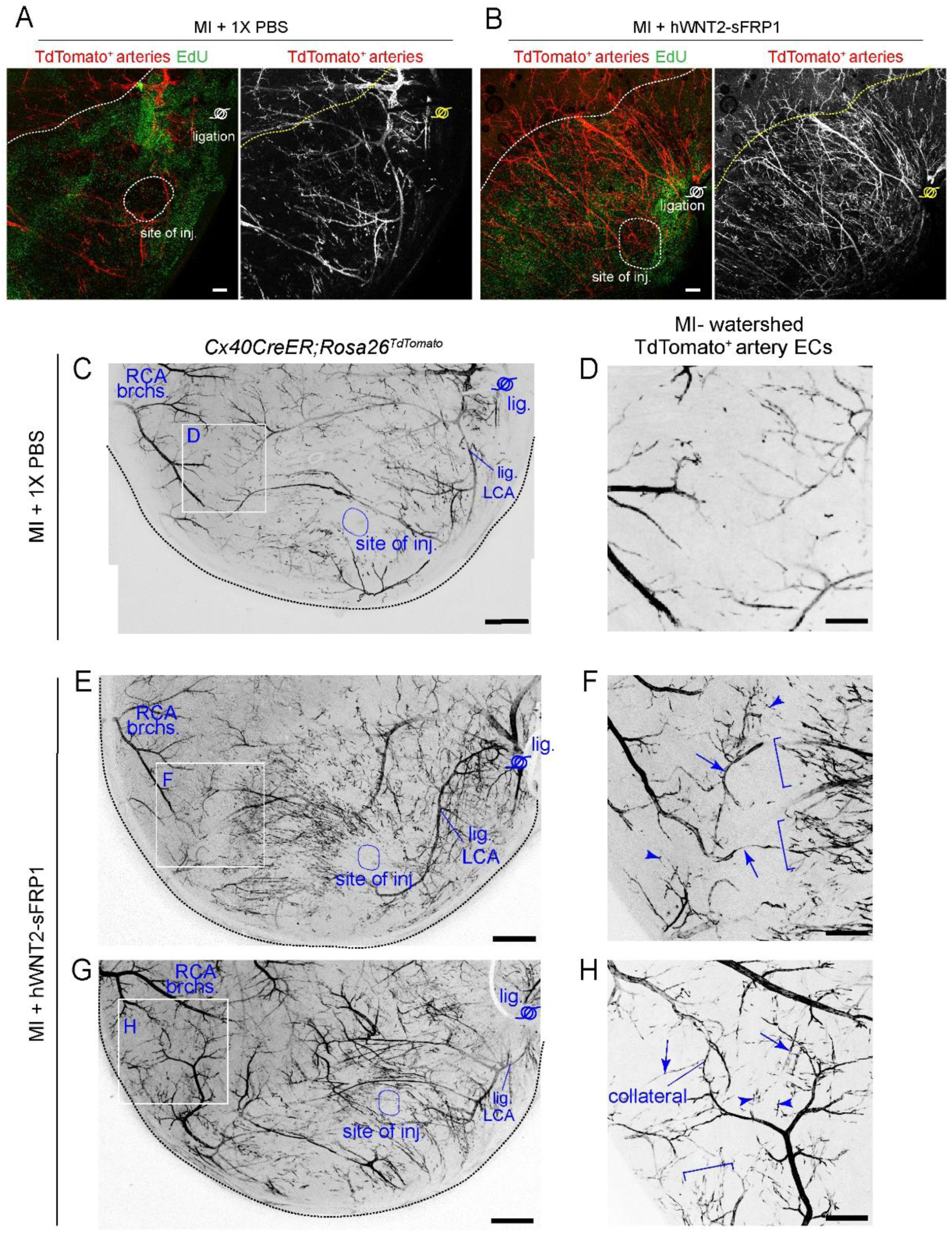
Exogeneous hWNT2-sFRP1 treatment in non-regenerative hearts. Experimental design is shown in Figure 1L. (**A**-**B**) Confocal image of P11 cardiac watershed area, injected with either (**A**) 1XPBS or (**B**) 55ng of hWNT2-sFRP1. TdTomato^+^ lineage traced artery endothelial cells are in red and EdU^+^ proliferating cells are in green. (**C**-**D**) Images of neonatal non-regenerative hearts injected with 1XPBS at the time of MI. **C** shows confocal image of stitched watershed region. **D** is an inset from **C**. (**E**-**H**) Images of neonatal non-regenerative hearts injected with 55ng of hWNT-sFRP1 at the time of MI. **E, G** show confocal images of stitched watershed regions. **F, H** are insets from **E** and **G** respectively. **F**, **H** show confocal images of 55ng of hWNT-sFRP1 injected watersheds. Arrowheads point to TdTomato^+^ single artery endothelial cells. Arrows point to TdTomato^+^ artery segments. Brackets mark clustering of TdTomato^+^ single artery endothelial cells. Scale bars: **A**, **B**, 500µm; **C**, **E**, **G**, 500µm; **D**, **F**, **H**, 200µm. MI, myocardial infarction; EC, endothelial cell; inj, injection; RCA, right coronary artery; brchs., branches; lig, ligation; LCA, left coronary artery; h, human; sFRP, soluble Frizzled receptor protein; Td, TdTomato.

**Supplementary Figure 6:**
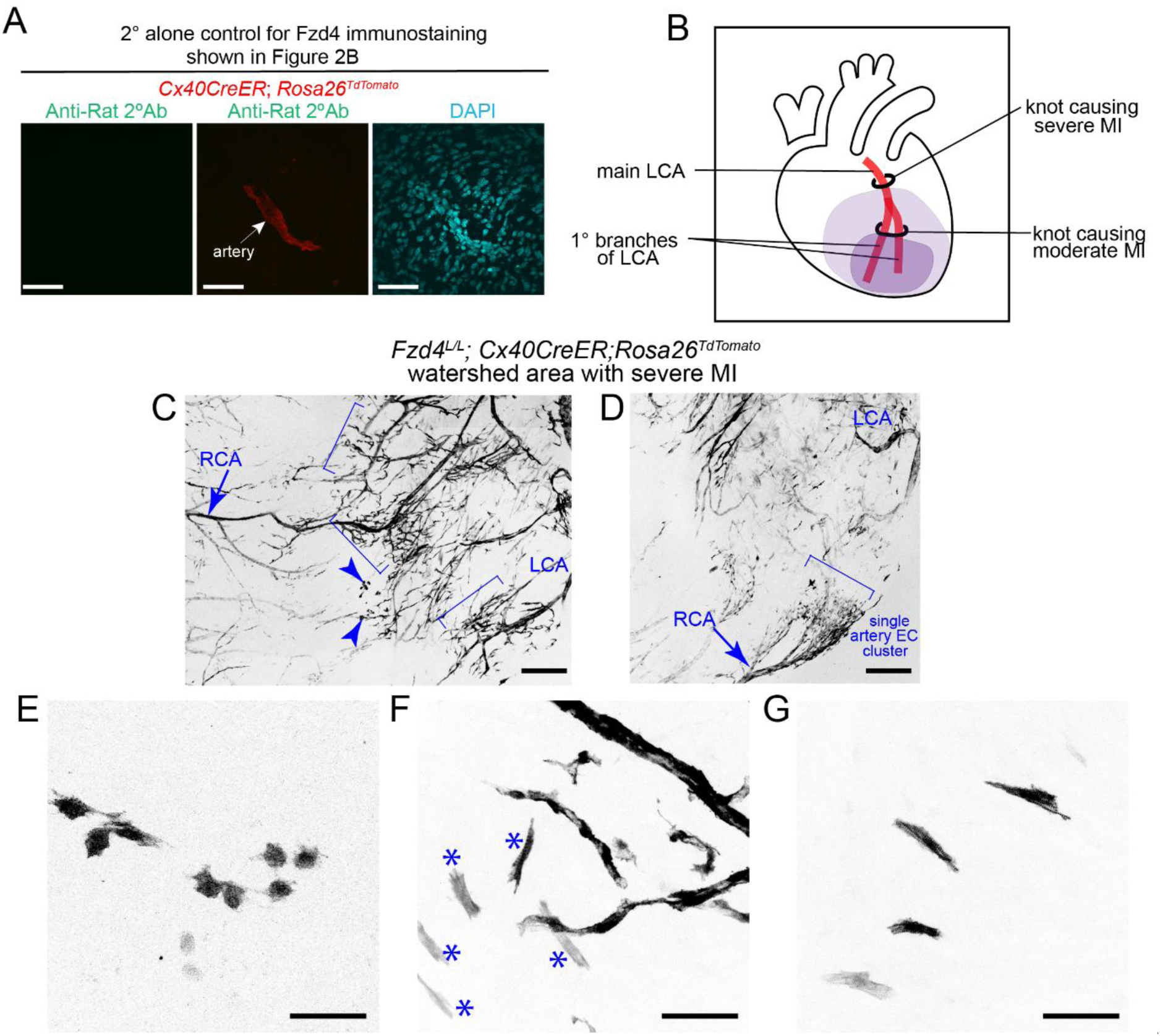
Arterial Fzd4 regulates endothelial morphologies. (**A**) Secondary alone control for immunostaining shown in Figure 2B. (**B**) Schematic showing positions of coronary occlusion through ligation of main LCA or primary branches of LCA to generate severe or moderate myocardial infarctions, respectively. (**C**, **D**) Confocal images of watershed regions from *Fzd4^L/L^; Cx40CreER; Rosa26^TdTomato^* hearts with severe MI. Arrowheads point to TdTomato^+^ lineage traced endothelial cells with non-EC morphologies. Brackets point to cluster of TdTomato^+^ endothelial cells. (**E**-**G**) Confocal images of TdTomato^+^ endothelial cells traced from pre-existing arteries showing non-EC morphology post-severe MI, in *Fzd4^L/L^; Cx40CreER; Rosa26^TdTomato^* hearts. * point to TdTomato-lineage traced cells with cardiomyocyte like features. Scale bars: **A**, **E**, **F**, **G**, 50µm; **C**, **D**, 200µm. 1°, primary; 2°, secondary; LCA, left coronary artery; RCA, right coronary artery; EC, endothelial cell; MI, myocardial infarction; Ab, Antibody.

**Supplementary Figure 7:**
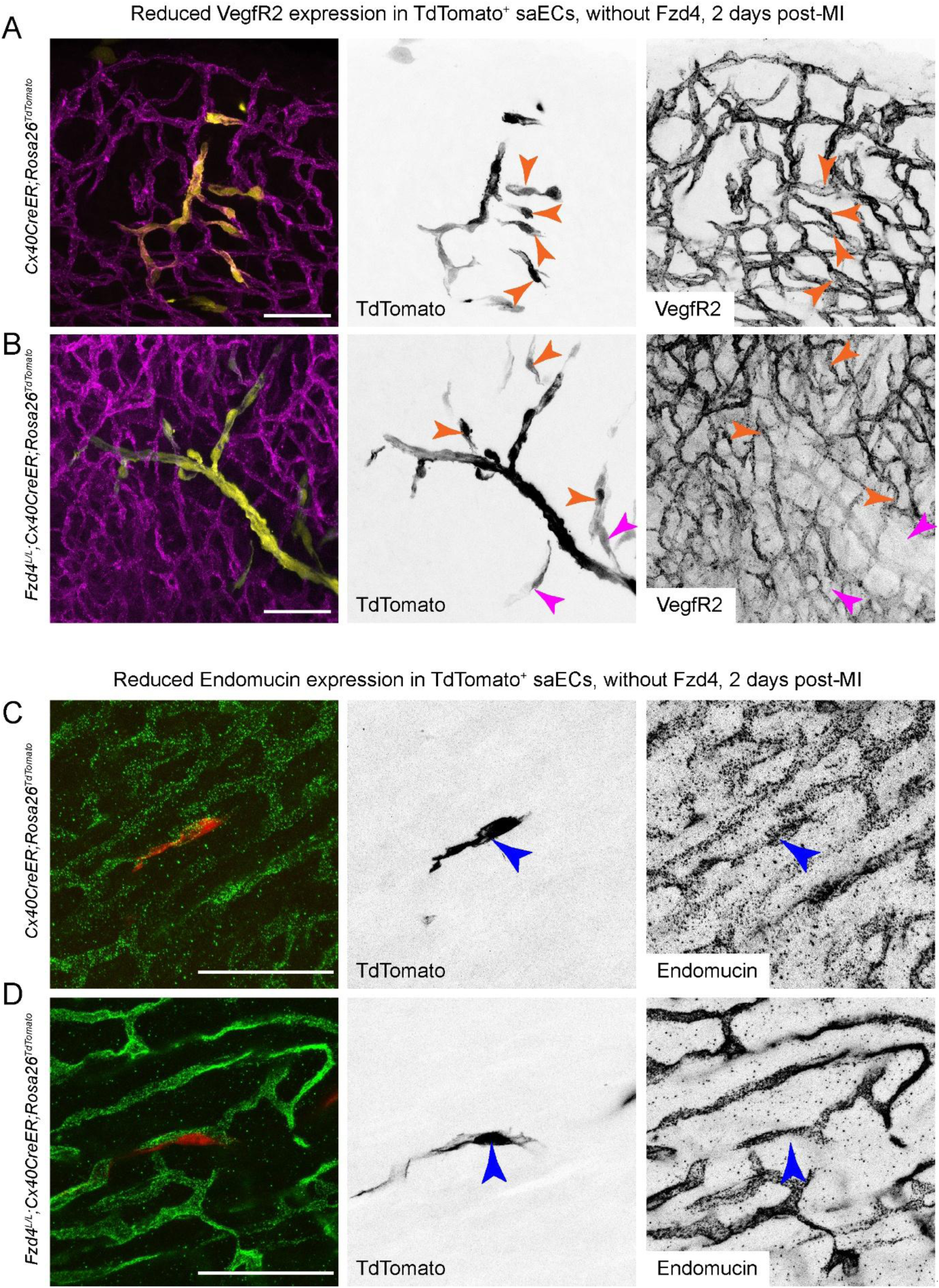
Fzd4 depleted artery ECs fail to express markers for de-differentiation. Experimental design is shown in Figure 3A. (**A**, **B**) Confocal images of watershed regions showing VegfR2 in magenta and single artery ECs in yellow, in (**A**) control and (**B**) *Fzd4* arterial knockout hearts. Orange arrowheads point to single cells with expression of VegfR2. Magenta arrowheads point to single artery cells with minimal expression of VegfR2. (**C**, **D**) Confocal images of watershed regions showing Endomucin in green and single artery ECs in red, in (**C**) control and (**D**) *Fzd4* arterial knockout hearts. Arrowheads point to single artery cells (**C**) with or (**D**) without Endomucin signal. Control: *Cx40CreER; Rosa26^TdTomato^*; Fzd4 arterial knockout: *Fzd4^L/L^; Cx40CreER; Rosa26^TdTomato^*. Scale bars: 50µm. saEC, single artery endothelial cell

**Supplementary Figure 8:**
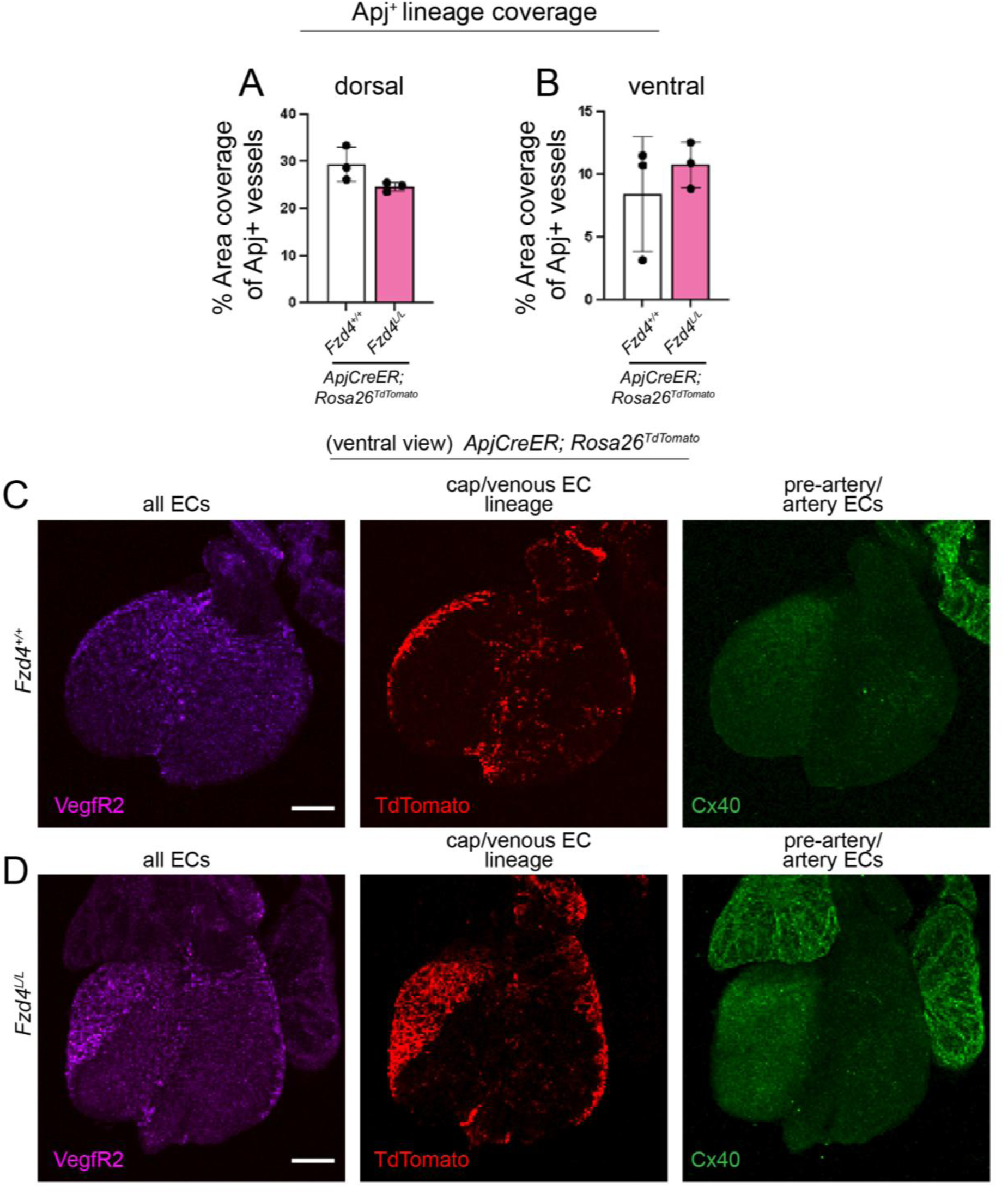
Absence of endothelial *Fzd4* does not affect capillary expansion during coronary artery development. (**A**, **B**) Quantification of area covered by Apj^+^ lineage cells in (**A**) dorsal and (**B**) ventral sides of developing hearts, at e13.5. Tamoxifen was injected at e10.5. p value=0.1459 (dorsal), p value=0.4866 (ventral) (**C**, **D**) Confocal images showing ventral view in hearts from (**C**) control and (**D**) *Fzd4* deleted from capillary endothelial cells. Magenta shows VegfR2, marks all endothelial cells. TdTomato marks Apj^+^ lineage cells. Cx40 marks artery endothelial cells. Scale bars: 200µm. cap, capillary; EC, endothelial cell; CT, control (*ApjCreER; Rosa26^TdTomato^*); knockout, (*Fzd4^L/L^; ApjCreER; Rosa26^TdTomato^*)

**Supplementary Figure 9:**
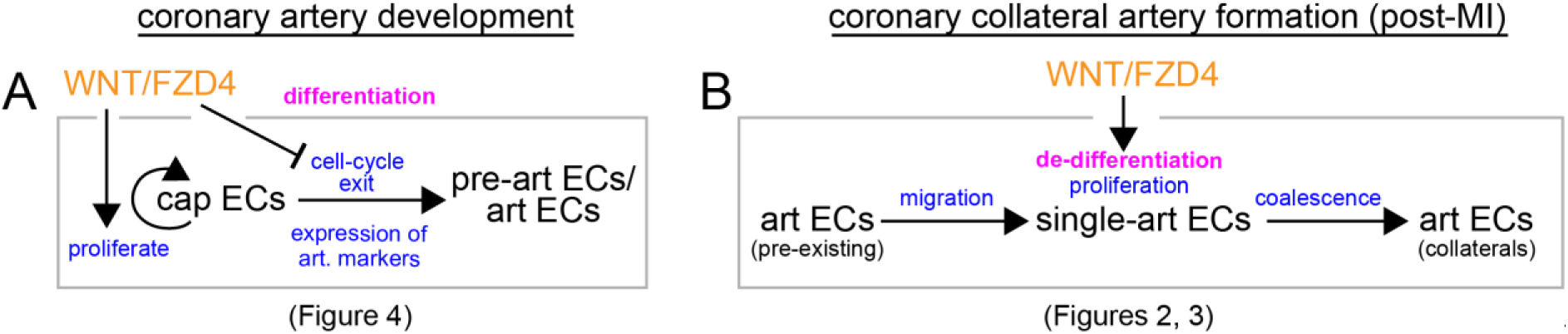
Role of endothelial Fzd4 in development of coronary arteries and post-MI collaterals. (**A**, **B**) Working model showing the potential role of endothelial Fzd4 in (**A**) development and (**B**) upon injury. (**A**) During coronary artery development, capillary Fzd4 is essential for its proliferative capacity and prevents capillary differentiation into arteries. In absence of capillary Fzd4, endothelial cells exit cell cycle, and differentiate into coronaries prematurely (supported by Figure 4). (**B**) Post-MI, arterial Fzd4 is essential for de-differentiation of single artery endothelial cells; absence of which, leads to reduction in expression of angiogenic/capillary markers (shown in Figure 3). Thus, arterial Fzd4 is required for coronary collateral formation by artery reassembly (shown in Figure 2).

**Supplementary Figure 10:**
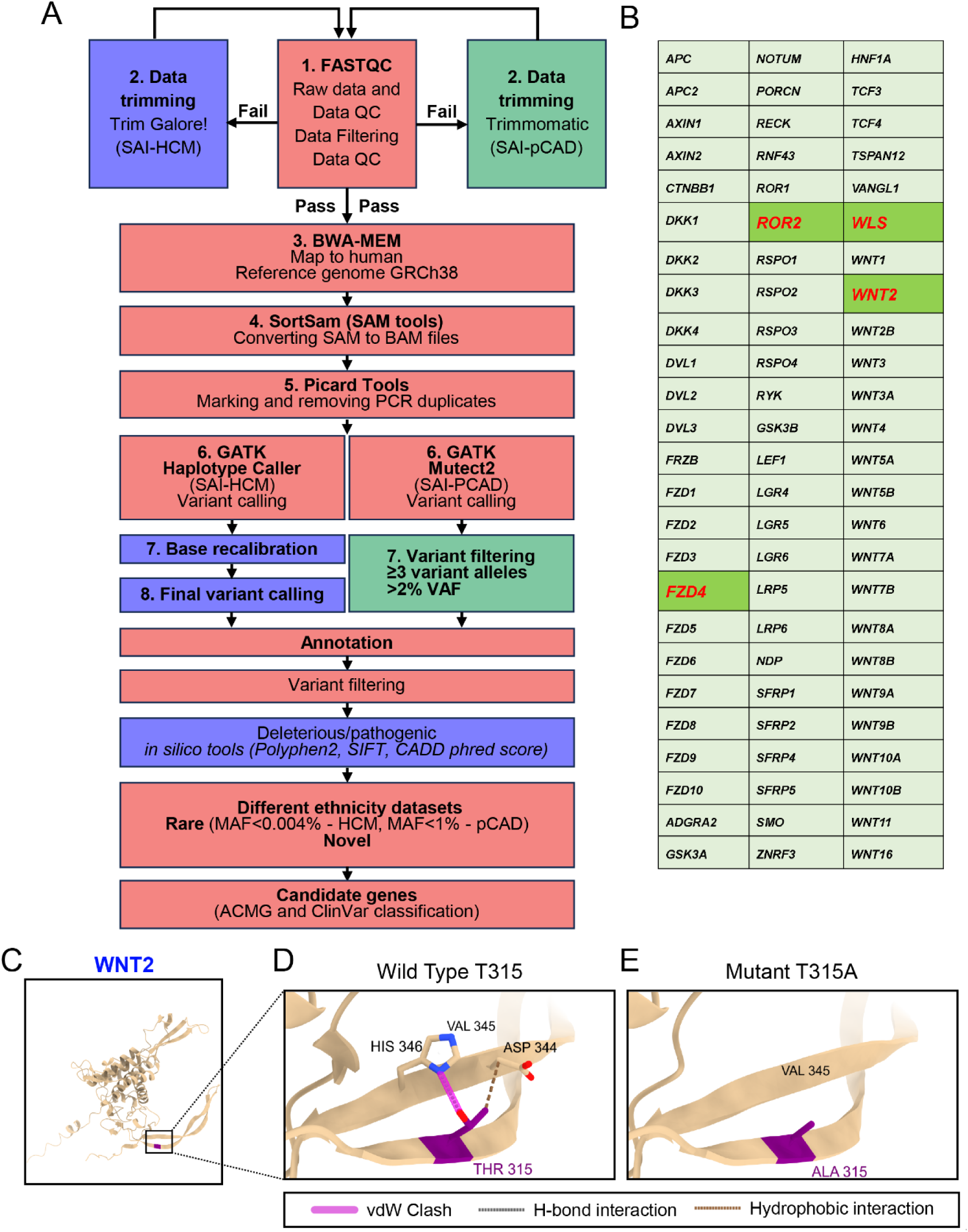
Genetic variants of WNT pathway genes in human heart patients. (**A**) Pipeline developed and acquired to analyse exome sequences from human patient samples. (**B**) List of 75 genes from WNT pathway checked for presence of genetic variants within the South Asian Indian population. Genetic variants of 4 genes (out of 75), namely, *WLS*, *WNT2*, *FZD4* and *ROR2* were determined to be variants of interest. (**C**-**E**) show predicted structural changes due to *WNT2* missense variant T315A. **(C)** shows the WNT2 structure obtained from the AlphaFold protein structure database. **(D, E)** show loss of H-bond (grey dotted line) and hydrophobic interactions (brown dotted line) in the T315A mutant could potentially disrupt the paired β strands in the WNT2 structure. ΔΔG value from DDmut for T315A mutation is 0.05 (stabilizing) with a missense score of 0.2213. SAI, South Asian Indian; HCM, Hypertrophic Cardiomyopathy; pCAD, premature Coronary Artery Disease; MAF, minor allele frequency; vdW, van der Waals

## Notes

### Competing Interest Statement

The authors have declared no competing interest.

